# A cheater founds the winning lineages during evolution of a novel metabolic pathway

**DOI:** 10.1101/2025.01.26.634942

**Authors:** Karl A. Widney, Lauren C. Phillips, Leo M. Rusch, Shelley D. Copley

## Abstract

Underground metabolic pathways—leaks in the metabolic network caused by promiscuous enzyme activities and non-enzymatic transformations—can provide the starting point for emergence of novel protopathways if a mutation or environmental change increases flux to a physiologically significant level. This early stage in the evolution of metabolic pathways is typically hidden from our view. We have evolved a novel protopathway in Δ*pdxB E. coli*, which lacks an enzyme required for synthesis of the essential cofactor pyridoxal 5′-phosphate (PLP). This protopathway is comprised of four steps catalyzed by promiscuous enzymes that are still serving their native functions. Complex population dynamics occurred during the evolution experiment. The dominant strain after 150 population doublings, JK1, had acquired four mutations. We constructed every intermediate between the Δ*pdxB* strain and JK1 and identified the order in which mutations arose in JK1 and the physiological effect of each. Three of the mutations together increased the PLP accumulation rate by 32-fold. The second mutation created a cheater that was less fit on its own but thrived in the population by scavenging nutrients released from the fragile parental cells. Notably, the dominant lineages at the end of the experiment all derived from this cheater strain.

---

Evolution of novel metabolic pathways has been a primary driver of organismal diversity on earth. Reconstructions of the genome of the last universal common ancestor (LUCA) suggest that core metabolic pathways had evolved by 3.8 billion years ago^1,2^. As ecosystems became more complex and new carbon sources became available, myriad degradative pathways evolved. Selective pressures to manipulate the environment and compete for resources fostered emergence of pathways for synthesis of complex natural products. Evolution of novel pathways continues in the present due to the introduction of anthropogenic chemicals, including pesticides^3–5^, pharmaceuticals^6^, solvents^7^, textile dyes^8^, explosives^9^ and myriad chemicals used in industry and household products.

Bioinformatic evidence suggests that metabolic pathways evolve by patching together enzymes that have a promiscuous ability to catalyze newly important reactions, followed by gene duplication and divergence to provide enzymes specialized for their new functions^10–14^. However, the retrospective view provided by bioinformatics reveals little about the *process* by which new pathways emerge. The suite of promiscuous activities available within a proteome and the mutations that allowed an initial “protopathway” to emerge are largely lost in time. We define “protopathways” as the earliest stage in the evolution of novel pathways. At this stage, promiscuous enzymes that have been recruited to serve new functions are still serving their native functions and proper regulation has not emerged. Protopathways likely follow the course of “underground pathways”^15^, minor leaks in the metabolic network caused by promiscuous enzyme activities and non-enzymatic decomposition reactions that normally do not impact fitness. Protopathways are different because a mutation or environmental change has made flux through the pathway physiologically relevant.

We have used adaptive laboratory evolution to probe the emergence of protopathways in bacteria in which the pathway for synthesis of the essential cofactor pyridoxal 5′-phosphate (PLP) has been broken by deletion of *pdxB* (Fig. 1). We previously evolved multiple lineages of Δ*pdxB E. coli* in glucose^16^. The Δ*pdxB* strain grew extremely slowly, but within 150 generations we obtained strains that grew at up to 74% the rate of wild-type *E. coli.* Strain JK1 restored PLP synthesis by patching together a novel protopathway (Fig. 1).

**Fig. 1.**
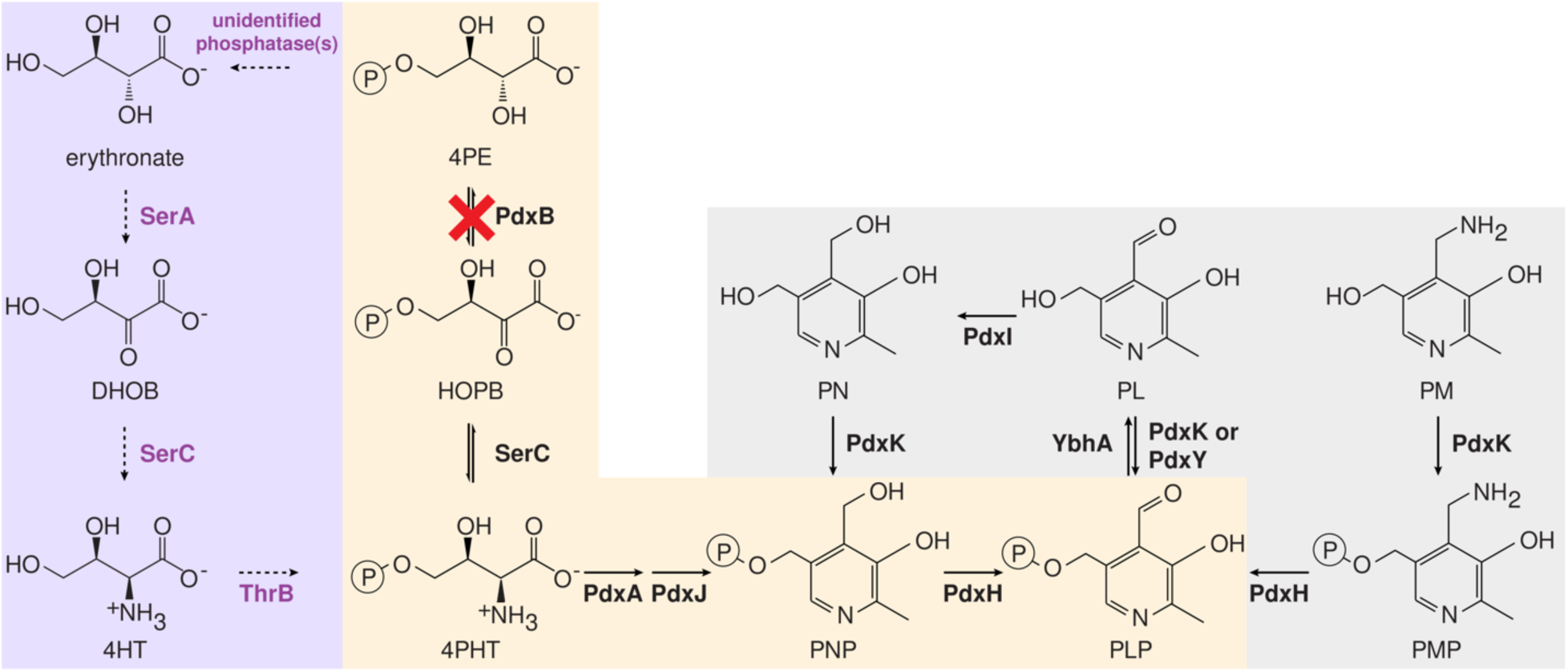
A four-step protopathway (highlighted in lilac) restores growth of Δ*pdxB E. coli.* Purple, promiscuous enzymes recruited to catalyze reactions in the protopathway; orange, canonical PLP synthesis pathway; grey, PLP vitamers and salvage enzymes. DHOB, (*3R*)-3,4-dihydroxy-2-oxobutanoate; 4HT, 4-hydroxy-L-threonine; 4PE, 4-phosphoerythronate; HOPB, (*3R*)-3-hydroxy-2-oxo-4-phosphooxybutanoate; 4PHT, 4-phosphooxy-L-threonine; PNP, pyridoxine 5′-phosphate; PLP, pyridoxal 5′-phosphate; PMP, pyridoxamine 5′-phosphate; PN, pyridoxine; PL, pyridoxal; PM, pyridoxamine, SerA, 3-phosphoglycerate dehydrogenase; SerC, phosphoserine/phosphohydroxythreonine transaminase; ThrB, homoserine kinase; PdxB, 4PE dehydrogenase; PdxA, 4PHT dehydrogenase; PdxJ, PNP synthase; PdxH, PNP/PMP oxidase; PdxK, PN/PL/PM kinase; YbhA, PLP phosphatase; PdxY, PL kinase.

We analyzed population genomic DNA at intervals during the evolution of JK1 to identify the sequence of mutations that led to JK1 as well as the overall population dynamics. Strain JK1 accumulated four mutations^16^: 1) a point mutation in *gapA* that decreases catalytic activity of glyceraldehyde 3-phosphate dehydrogenase; 2) a frameshift mutation in *rpoS*, which encodes the master regulator of the general stress response (σ^S^); 3) a large deletion that destroys the broad-specificity phosphatase YbhA and the pentose phosphate pathway enzyme Pgl (6-phosphogluconlactonase); and 4) a point mutation in *rpoC*, which encodes the β′ subunit of RNA polymerase.

We defined the fitness landscape for evolution of JK1 by reconstructing all intermediates between Δ*pdxB E. coli* and strain JK1 and characterizing their rates of growth and PLP accumulation. Multiple trajectories could have led to JK1. Interestingly, the actual trajectory passed through an intermediate that grows quite poorly on its own. Growth of this intermediate was stimulated by unidentified compounds released into the medium by its progenitor, demonstrating that modification of the environment by clones in an evolving population can significantly impact population dynamics. The initial *gapA\** mutation and the later deletion of *ybhA* contributed to improved PLP levels. The *rpoS\** mutation allowed the *gapA* rpoS\** strain to cheat off other cells in the population. The final point mutation in *rpoC* improves growth in minimal medium, a finding consistent with results of previous studies^17–19^.

## Results

### Complex population dynamics preceded the emergence of JK1

We sequenced genomic DNA from archived samples collected during the evolution of JK1 to identify when mutations in dominant lineages occurred. Clones with different point mutations in *gapA,* which encodes glyceraldehyde 3-phosphate dehydrogenase (GAPDH), arose early (Fig. 2). The *k_cat_/K_M_* of G210V GAPDH is eight-fold lower than that of the wild-type enzyme. L158Q GAPDH has normal catalytic activity but is less stable than the wild-type enzyme; its T_M_ is decreased from 72.6 ± 0.2 °C to 66.5 ± 0.4 °C. These mutations decrease GAPDH activity in lysates by 60-80%^16^. A third mutation in *gapA* changes Asp34 to Asn. Asp34 coordinates the hydroxyl groups of the NAD(H) cofactor (PDB 1GAD); changing this residue to Asn should impair NAD(H). binding.

**Fig. 2.**
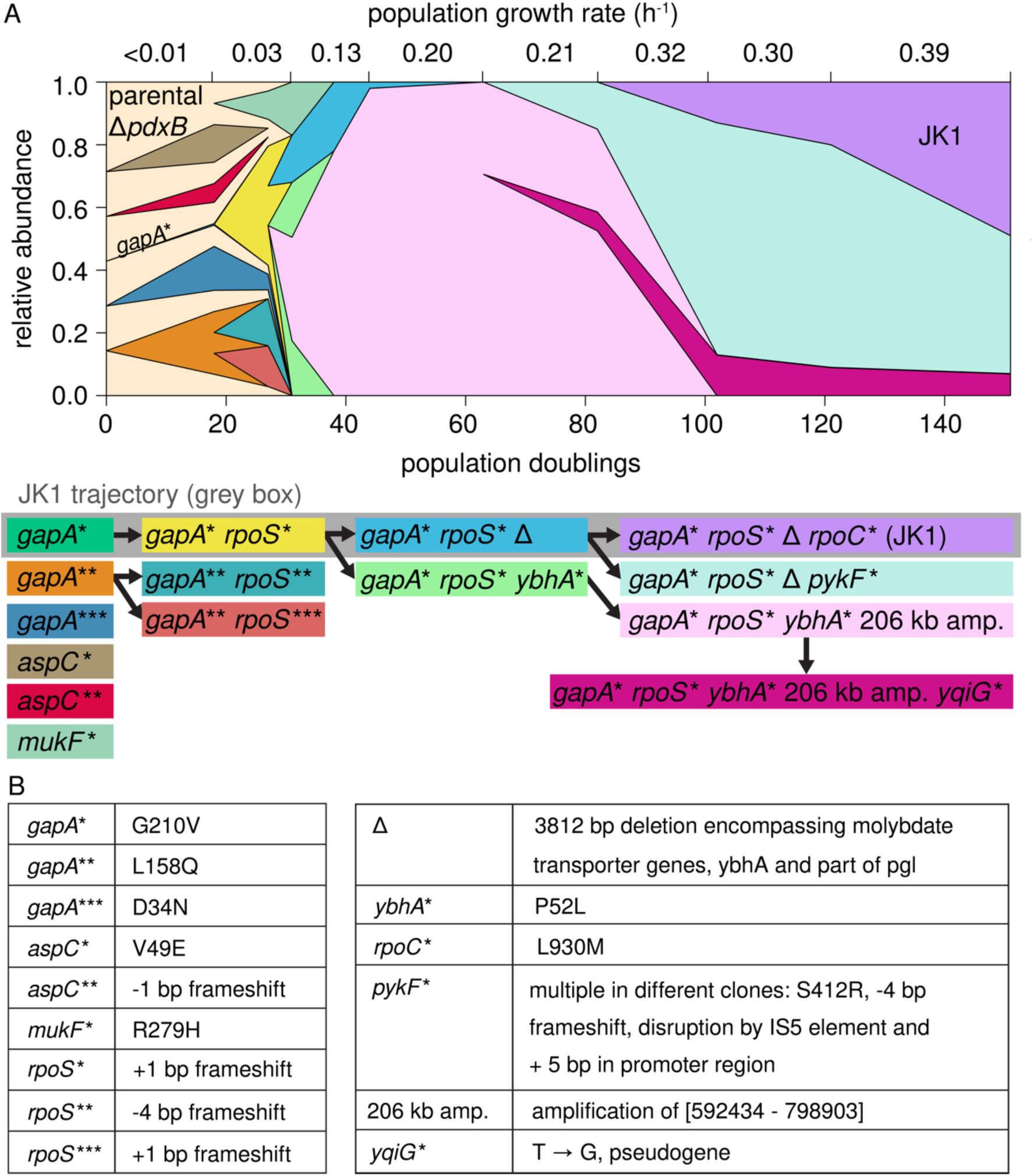
Complex population dynamics occurred during evolution of strain JK1. (A) Muller plot tracking abundance of clones throughout the evolution experiment. To simplify the analysis, the plot tracks only mutations that were found at two or more consecutive timepoints, were present in over 10% of the reads or arose in a gene in which other mutations had already been found. (B) Mutations listed in A. Additional details are provided in Supplementary Data Set 1.

Clones with mutations in *gapA* quickly acquired different frameshift mutations in *rpoS* that likely abrogate function. One *rpoS* mutation arose in the *gapA\** background in the lineage that eventually led to JK1. Two other *rpoS* mutations arose in the *gapA\*\** background. At 25 generations, strains with mutations in *gapA* and *rpoS* comprised 68% of the population, suggesting that this combination provides a significant improvement in fitness.

Two clones emerged from the *gapA* rpoS\** clone. The clone that led to JK1 acquired a large deletion encompassing the molybdate transport operon, *ybhA* and part of *pgl* (cyan clone in Fig. 2). (This deletion will subsequently be referred to as Δ for simplicity.) Another clone acquired a point mutation in *ybhA* (*ybhA*\*, light green clone) and then amplified a 206 kb region encompassing *ybhA\** (Fig. E1, pink clone in Fig. 2). This clone overtook the population within about 10 generations. The clone leading to JK1 was not detectable at 63 population doublings. Given the 92x sequence coverage, the *gapA* rpoS\** Δ clone must have comprised less than 1% of the population at this point. However, additional mutations in either *rpoC* (lilac clone) or *pykF* (turquoise clone) enabled the *gapA* rpoS\** Δ lineage to overcome the previously dominant lineage, possibly due to the burden imposed by the amplified 206 kb region.

### Impaired PLP synthesis compromises the integrity of the cell wall

Synthesis of peptidoglycan requires substantial quantities of L-alanine, D-alanine, D-glutamate and meso-diaminopimelic acid. Three of these compounds are produced by PLP-dependent enzymes. Not surprisingly, the *ΔpdxB* and *gapA*\* strains show signs of a weak cell wall. These strains have abnormal morphologies (Fig. 3). Further, elevated levels of DNA and B_6_ vitamers (including PLP as well as pyridoxal (PL), pyridoxine (PN), pyridoxamine (PM), PNP and PMP, Fig. 1) are found in spent medium after growth of these strains. Impaired PLP synthesis apparently weakens the cell walls of these strains, leading to lysis of some cells with release of cytoplasmic contents into the medium. Oddly, the *gapA* rpoS\** Δ strain formed long filaments, suggesting a perturbed balance between cell elongation and cell division. In contrast, the evolved JK1 cells have normal morphology and release much less DNA and vitamin B_6_ into the spent medium than the parental Δ*pdxB* and the *gapA\** strains.

**Fig. 3.**
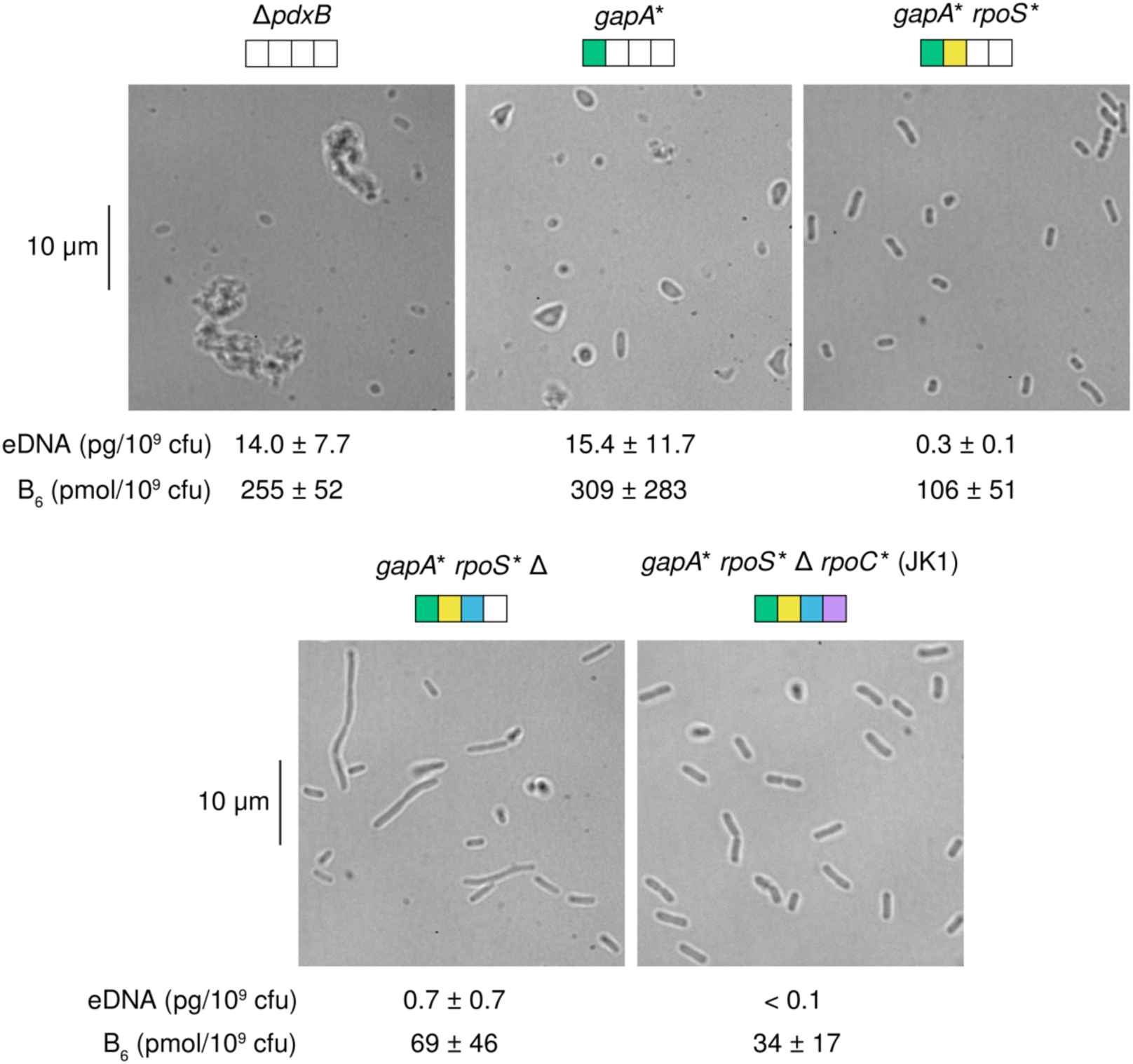
The parental Δ*pdxB E. coli* and intermediate strains exhibit odd cell morphologies. Amounts of extracellular DNA (eDNA) and B_6_ vitamers in the spent medium are provided below each micrograph.

### Most evolutionary trajectories toward JK1 are accessible

Strain JK1 had accumulated four mutations in the order depicted in Fig. 2. To determine whether other evolutionary trajectories were accessible, we constructed all possible intermediates between Δ*pdxB E. coli* and JK1 and measured their growth rates in M9/glucose (Fig. 4, top numbers in each circle). The resulting fitness landscape reveals several interesting features. First, the *gapA*\* mutation was the only one of the four mutations that increased the growth rate of the parental Δ*pdxB* strain. Second, the *rpoS\** mutation was not beneficial in any background and, indeed, reduced the growth rate of the *gapA\** strain by 4-fold. This finding is unexpected, since the Muller diagram in Fig. 2 clearly shows that *rpoS* mutations were advantageous in the background of a *gapA* mutation. We will return to this puzzling finding below. Third, the Δ mutation only improved growth rate when the *gapA* mutation was already present. Finally, the *rpoC* mutation was only beneficial when the *gapA\** and Δ mutations were already present.

**Fig. 4.**
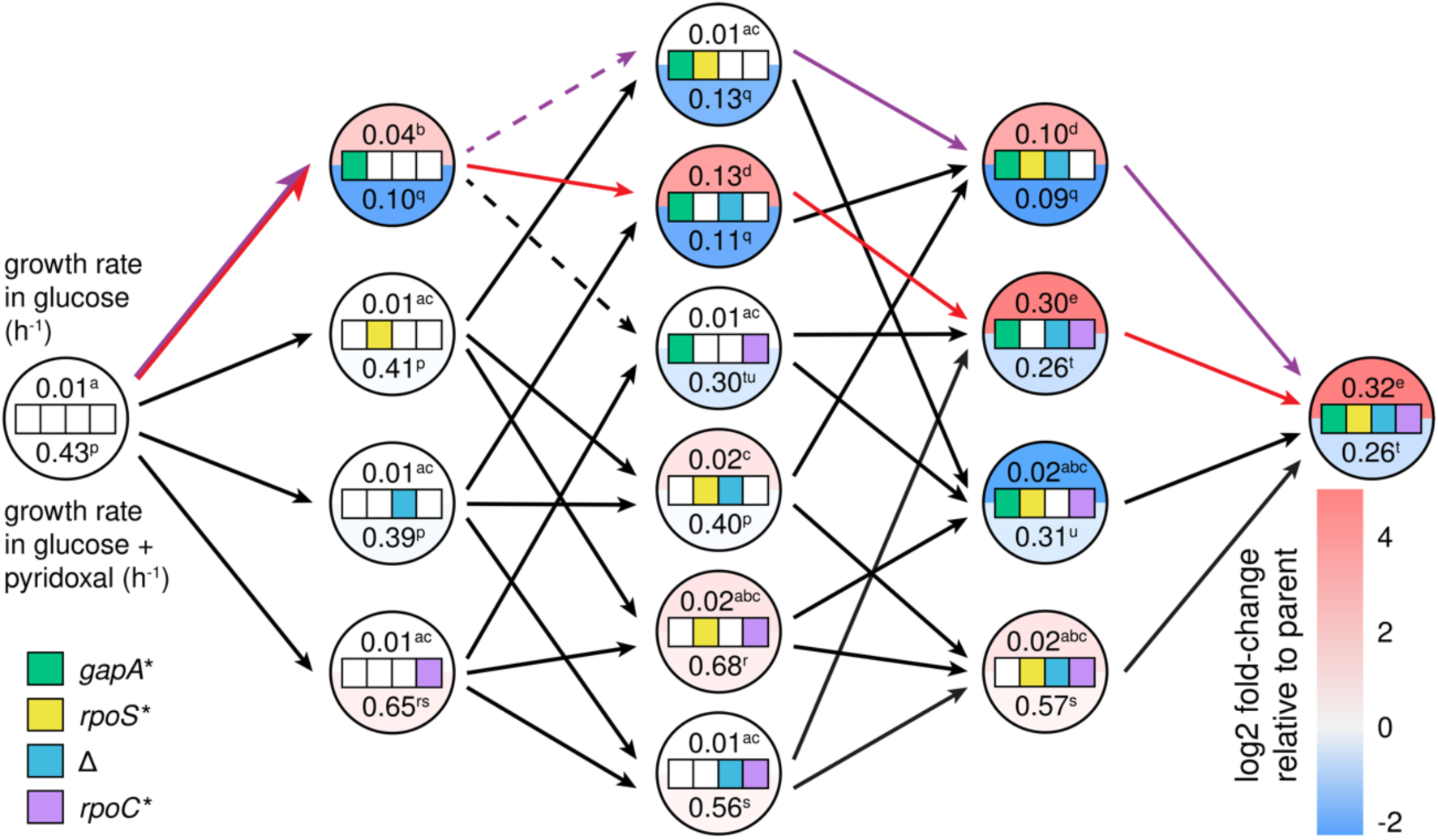
Fitness landscape showing the effects of each mutation on growth in M9/glucose in the absence (top) and presence (bottom) of 10 µM pyridoxal, which can be converted to PLP by a salvage enzyme (Fig. 1). Superscripts represent statistical groups with p-adj < 0.05 after Dunnett T3 correction for multiple comparisons (Graphpad Prism 8). Purple arrows, the actual trajectory by which JK1 evolved; red arrows, the optimal trajectory. Dashed lines indicate steps that result in a statistically significant decrease in growth rate in M9/glucose.

The fitness landscape in M9/glucose (Fig. 4, top numbers in each circle) reveals positive epistasis between the mutations that led to JK1. In the absence of epistasis, we would expect JK1 to grow four-fold faster than Δ*pdxB E. coli*; the actual increase in growth rate is 32-fold. The *rpoC\**, *rpoS\** or Δ mutations have little or no effect in most backgrounds, creating a smooth fitness landscape with 30 of 32 mutational steps accessible and all 16 intermediates accessible by at least one step. Only one trajectory (red arrows in Fig. 4) had significant increases in growth rate at each of the first three steps. Notably, the actual trajectory toward JK1 proceeded through one of only two mutational steps (dashed arrows in Fig. 4) that significantly reduced fitness.

### The *gapA\** mutation is the critical first step

The *gapA\** mutation is clearly the critical first step toward establishing the new protopathway. The *gapA*\* mutation occurred first in the lineage leading to JK1. Two other clones also acquired mutations in *gapA* prior to other mutations. Clones that acquired mutations in other genes (*aspC* and *mukF*) were quickly outcompeted in the evolving population (Fig. 2).

The *gapA\** mutation increased growth rate in M9/glucose by four-fold in the parental background and up to thirty-fold in other backgrounds (e.g. Δ *rpoC\** ® Δ *rpoC* gapA\**) (Fig. 2). To determine whether this increase in growth rate was due to an increase in PLP synthesis, we optimized a published procedure for measuring B_6_ vitamers^20^. We measured the total level of B_6_ vitamers (Fig. 1) in cell lysates and spent medium to calculate the total rate of B_6_ accumulation. Assuming that the total B_6_ content must be doubled each time the cells divide, we can calculate the rate of PLP accumulation by dividing total B_6_ content by the doubling time. We term this an accumulation rate because B_6_ content is the net result of its synthesis minus its degradation. Obtaining accurate B_6_ levels for the parental strain was difficult because it was so fragile and prone to lysis (Fig 3). However, the growth rates of the Δ, *rpoS*\* and *rpoC*\* strains are similar to that of the parental strain, suggesting that the parental strain likely has a similar PLP accumulation rate, approximately 3 pmol/(10^9^ cfu)(h) (Fig. 5). The *gapA*\* strain has a PLP accumulation rate of 9 pmol/(10^9^ cfu)(h) (Fig. 5), suggesting that the *gapA*\* mutation increases PLP accumulation rate by about three-fold, a value commensurate with the four-fold increase in growth rate.

**Fig. 5.**
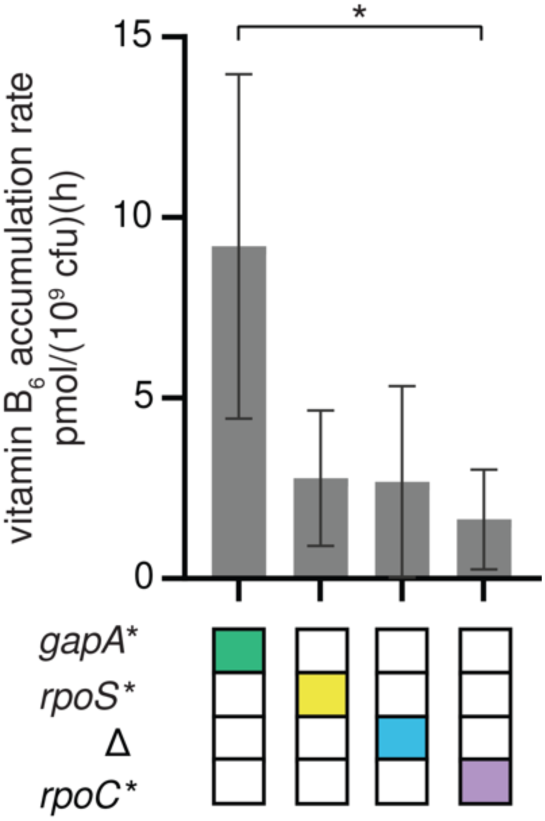
PLP accumulation rates in single mutants suggest that the *gapA\** mutation increases PLP synthesis. Error bars represent 95% confidence intervals. P-values adjusted by Dunnett T3 correction for multiple comparisons (Graphpad Prism 8). *, p-adj < 0.05.

The *gapA\** mutation likely improves flux through the protopathway by alleviating the feedback inhibition of SerA, the first enzyme in the serine synthesis pathway, by serine. Previous work showed that the *gapA\** mutation reduces the activity of GapA in cell lysates by 80%^16^. This bottleneck in glycolysis likely decreases the concentration of downstream metabolites. One of these, 3-phosphoglycerate, is the native substrate of SerA. A reduction in the level of 3-phosphoglycerate should decrease production of serine, thereby reducing allosteric inhibition of SerA by serine. Previous metabolomics data showed that the mutations in JK1 reduce serine level below the limit of detection^16^. The improvement in PLP synthesis due to the *gapA\** mutation suggests that it plays a major role in reducing serine levels.

Although the *gapA\** mutation is critical for increasing flux through the protopathway, it comes with a clear cost. Adding the *gapA\** mutation to every background decreases growth rate in non-selective conditions (M9/glucose plus pyridoxal) (Fig. 4, bottom values in circles). Trade-offs caused by mutations that are beneficial under selective conditions but detrimental once selective pressures are relieved are common^21,22^. We refer to such mutations as “expedient” to denote their ability to quickly solve a problem under one condition, but at an overall fitness cost. The *gapA\** mutation is clearly expedient.

### The *gapA* rpoS\** intermediate strain in the trajectory toward JK1 is a cheater

Remarkably, JK1 evolved by a trajectory that passes through an intermediate whose growth rate in M9/glucose is slower than that of its progenitor; addition of *rpoS\** to the *gapA\** background results in a significant decrease in growth rate (Fig. 4). Yet, the *gapA* rpoS\** clone outcompeted its progenitor, the *gapA\** clone (Fig. 2). Additionally, two other *rpoS* mutations arose in a clone with a different *gapA* mutation (teal and terracotta clones in Fig. 2). These results suggest that, counter to expectations based upon the fitness landscape, loss of function mutations in *rpoS* were beneficial in the evolving population during the evolution of Δ*pdxB E. coli*.

Mutations in *rpoS* are common in experimental evolution in M9/glucose^23–26^. If the *rpoS\** mutation were providing a general growth benefit, we would expect it to increase growth rate when selection for improved PLP synthesis is alleviated by addition of pyridoxal to the medium. However, the *rpoS\** mutation did not increase growth rate in M9/glucose plus pyridoxal in any background (Fig. 4). The lack of a beneficial effect of the *rpoS\** mutation when clones are grown in isolation suggests that the *gapA* rpoS\** strain is a cheater that takes advantage of resources provided by other clones in the evolving population. This notion is consistent with a recent report that loss-of-function mutations in *P. putida rpoS* allowed strains to cheat off other strains with intact *rpoS*^27^. To test this hypothesis, we measured the growth rates of the *gapA\** and *gapA* rpoS\** strains in spent medium collected after growth of the parental cells (Fig. E2). The *gapA* rpoS\** strain showed a substantial improvement in growth rate, suggesting that this strain exploits resources released into the medium by lysis of the fragile parental Δ*pdxB* cells.

The slow growth rate of the *gapA* rpoS\** clone in M9/glucose compared to that of its progenitor suggests that the RpoS stress response is important for growth of the *gapA\** clone in M9/glucose. Consistent with this hypothesis, the *gapA*\* mutation results in upregulation of 39 genes in the RpoS regulon compared to the parental Δ*pdxB* strain (Fig. E3, Supplementary Data Set 2). Several of the upregulated genes encode glycolytic enzymes, including glucose 6-phosphate isomerase, fructose bisphosphatases A and B, GAPDH and enolase. Thus, we suspect that activation of the stress response in the *gapA*\* clone helps compensate for the decrease in glycolytic flux caused by the *gapA*\* mutation. However, loss of RpoS in the *gapA* rpoS\** clone apparently allows this strain to avoid activating the programmed slowdown in growth regulated by RpoS. Thus, this strain is a cheater, taking advantage of resources released from other cells in the population but providing nothing in return. Interestingly, every successful clone by the end of the experiment originated from the cheater.

### The Δ mutation improves PLP accumulation rate in backgrounds containing the *gapA*\* mutation

The addition of Δ to the *gapA*\* *rpoS\** clone causes a 10-fold-increase in growth rate and a 32-fold increase in PLP accumulation rate (Fig. 6). The Δ mutation also increases growth rate in other *gapA*-*containing backgrounds, ranging from 3.3-fold (e.g. *gapA\** ® *gapA\** Δ) to 30-fold (e.g. *gapA* rpoC\** ® *gapA* rpoC\** Δ) (Fig. 4). The addition of Δ to backgrounds lacking the *gapA\** mutation results in at most a two-fold increase in growth rate, suggesting that *gapA\** is required to unlock the benefits of Δ.

**Fig. 6.**
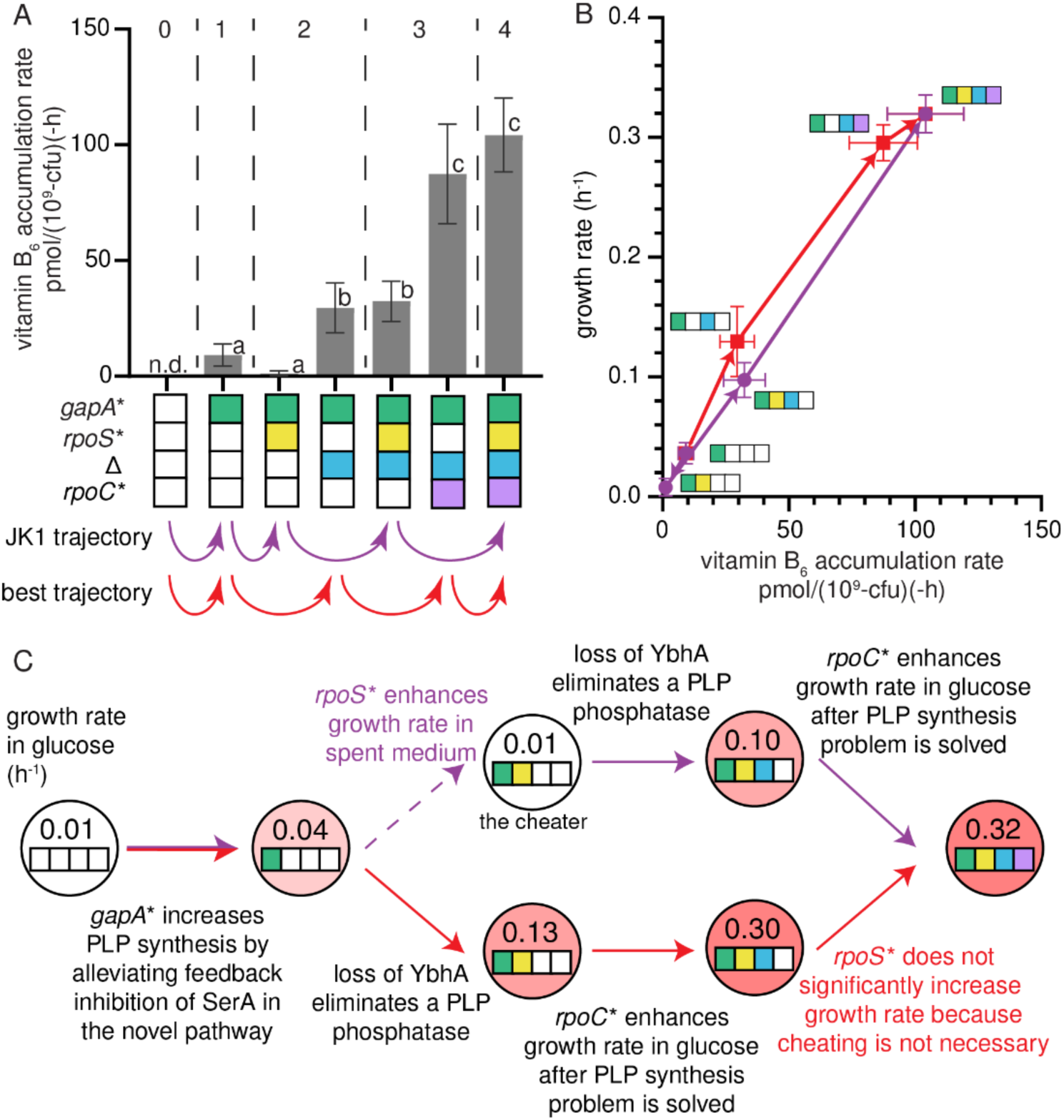
The combination of the *gapA\** and Δ mutations significantly increases PLP accumulation rate. A) Vitamin B_6_ accumulation rates in the actual and best trajectories. Grey dashed lines separate the parent, single, double, triple and quadruple mutant strains. Error bars represent 95% confidence intervals. Letters indicate statistical groups with p-adj < 0.05 after Dunnett T3 correction for multiple comparisons (GraphPad Prism 8). n.d., not done. B) Growth rate versus vitamin B_6_ accumulation rate. Arrows indicate evolutionary trajectories. Error bars represent 95% confidence intervals. C) Summary of the mutations and their effects in the trajectory leading to JK1 and the best trajectory.

The Δ mutation removes two genes of interest, *ybhA* and *pgl*. Loss of YbhA, a PLP phosphatase^28^, should spare PLP from degradation but would be of little help unless a prior mutation had increased flux through the protopathway. Parental Δ*pdxB* cells and others lacking the *gapA\** mutation likely have extremely low levels of free PLP, so YbhA would have little substrate in these cells. However, after the *gapA\** mutation increases flux through the protopathway, levels of PLP are apparently high enough for loss of YbhA to be beneficial. Increasing intracellular PLP levels resulting from the *gapA\** mutation and the deletion of *ybhA* could create a positive feedback loop that increases flux through the protopathway. SerC, which catalyzes the transamination of DHOB in the protopathway (Fig. 1), requires PLP. Thus, increasing PLP may increase the concentration of active SerC, further enhancing production of PLP.

We previously hypothesized that removal of *pgl* would result in diversion of GAP from glycolysis into the pentose phosphate pathway, thereby decreasing flux into the lower glycolytic pathway and contributing to the decrease in serine concentration that relieves feedback inhibition of SerA^16^. However, a clone of *gapA* rpoS\** that acquired a point mutation in *ybhA* (with no effect on *pgl*) significantly increased in abundance in the population (Fig. 2), suggesting that the *ybhA* mutation alone is sufficient to cause a significant increase in growth rate.

The *gapA* rpoS\** Δ clone is the first intermediate in the trajectory to JK1 that has similar growth rates in M9/glucose and M9/glucose plus pyridoxal (Fig. 4), suggesting that its growth is not limited by PLP synthesis. However, the physiology of this strain is still clearly abnormal because the cells form long filaments (Fig. 2).

### The *rpoC\** mutation substantially improves growth rate in glucose

The addition of *rpoC\** to the *gapA* rpoS\** Δ strain results in a 3-fold increase in growth rate in M9/glucose (Fig. 4) and a similar increase in M9/glucose plus pyridoxal. *rpoC\** increases growth rate in M9/glucose in one other strain, *gapA\** Δ, whose growth also does not appear to be limited by PLP accumulation. Thus, *rpoC\** confers an increase in growth rate only after PLP accumulation has been restored, suggesting that it simply improves growth rate in glucose. In agreement with this finding, the addition of *rpoC\** to all backgrounds improves growth rate in M9/glucose plus pyridoxal. Additionally, mutations in *rpoC* are frequently observed after adaptive laboratory evolution in glucose and glycerol and have many effects that improve growth rate in minimal medium^18,19,29^.

Interestingly, the rate of PLP accumulation increases after the *rpoC*\* mutation. The comparable growth rates of the *gapA* rpoS\** Δ strain and JK1 in the absence and presence of pyridoxal suggest that the ability to synthesize PLP no longer limits growth rate in either strain. Apparently flux through the protopathway can be increased to keep up with the demand required by the faster growth conferred by the *rpoC*\* mutation. However, we cannot rule out the possibility that some of the transcriptional changes caused by the *rpoC*\* mutation play a role in increasing PLP synthesis. *rpoC* mutations are known to increase metabolic efficiency and anabolism during growth in minimal media and have pleiotropic effects on the transcriptome^18^, proteome and metabolome^17^. For example, a mutation in *rpoC* perturbs the levels of 68 metabolites and 118 proteins when *E. coli* MG1655 is grown in glycerol as a sole carbon source^17^. Thus, it is difficult to determine whether the *rpoC\** mutation directly increased PLP accumulation rate in JK1.

## Discussion

By characterizing the fitness landscape for the evolution of JK1, we identified how mutations lead to the emergence of a novel protopathway (Fig. 6C, purple arrows). The *gapA\** mutation increases the rate of PLP synthesis by reducing feedback inhibition of SerA, allowing it to better perform its new function in the protopathway. The next mutation, *rpoS\**, allows the *gapA* rpoS\** strain to cheat off other strains in the population. Since the stress response activated by RpoS shifts metabolism toward slow growth when the cells perceive stress or starvation^26,30^, loss of RpoS apparently allows the *gapA* rpoS\** strain to grow more rapidly using the abundant glucose in the medium as well as the metabolites released from lysis of the fragile parental Δ*pdxB* cells. The Δ mutation is only beneficial in backgrounds containing the *gapA\** mutation, suggesting that loss of the PLP phosphatase YbhA is only beneficial after PLP synthesis has been improved by the *gapA\** mutation. Finally, the *rpoC\** mutation increases growth rate in glucose only after PLP accumulation rate no longer limits growth rate.

The “best” trajectory toward JK1 is the one with the largest fitness increase at each step. This trajectory (Fig. 6C, red arrows) also begins with the *gapA\** mutation. However, this trajectory does not pass through the *gapA* rpoS\** cheater strain. Bypassing the cheater mutation enables the clone following the best trajectory to reach a growth rate comparable to that of JK1 in M9/glucose by only three mutations. If JK1 had been characterized solely by reverting individual mutations, *rpoS\** might have been mistakenly identified as a hitchhiker mutation (i.e. a mutation that confers no growth benefit but became fixed during evolution because it coincided with a beneficial mutation). However, its evolutionary significance is evident, as the *gapA* rpoS\** clone became a dominant strain in the population (Fig. 2) by exploiting other strains.

All clones in the population by the end of the evolution depicted in Fig. 2 arose from the *gapA* rpoS\** cheater strain; JK1 was the dominant clone. Several other successful clones (grouped together in turquoise in Fig. 2) also arose in the *gapA* rpoS\** Δ background by acquiring mutations in *pykF.* Mutations in *pykF* were also found in two other evolved populations of ***Δ****pdxB E. coli*^16^. Mutations in *pykF* have been identified after long-term evolution of *E. coli* in glucose^31^. Thus, the occurrence of multiple *pykF* mutations and the *rpoC\** mutation in the *gapA* rpoS\** Δ background is consistent with the conclusion that PLP accumulation rate is adequate at this stage and selection has shifted to favor mutants that grow faster in minimal medium containing glucose as a sole carbon source.

Although the mutations in JK1 greatly increase growth rate in M9/glucose compared to the parental Δ*pdxB* strain, JK1 grows at only 50% of the rate of wild-type *E. coli* in M9/glucose^16^. Previous work showed that the concentrations of 3-phosphoglycerate, a glycolytic intermediate, and serine, which is synthesized from 3-phosphoglycerate, are very low in JK1^16^. Thus, the mutations that improved PLP accumulation rate in JK1 caused significant perturbations to the metabolic network. In theory, the need for the damaging *gapA\** mutation could be alleviated by duplication and divergence of *serA* to evolve an efficient erythronate dehydrogenase that is not subject to inhibition by serine. That said, this protopathway is not an elegant solution to the problem of restoring PLP synthesis because it wastes ATP; a phosphate is removed in the first step of the protopathway and restored in the last step. Thus, the most accessible protopathway that arises in Δ*pdxB E. coli* may not be the best solution in the long run.

## Online Methods

### Biological resources

Strains, plasmids and primers used in this work are included in Tables S1, S2 and S3, respectively, in Supplementary Information. Oligonucleotides and gene blocks (gBlocks) used to construct guide and donor plasmids are included in Tables S4 and S5, respectively. gBlocks used to construct protein expression plasmids are included in Table S6.

### Statistical analyses

Unless otherwise noted, statistical analyses were performed in Graphpad Prism 8. Statistical groups were assigned in R using the multcompletters function.

### Reagents

Chemicals (antibiotics, IPTG, etc.) and media (LB and SOC) were purchased from Sigma-Aldrich and Thermo-Fischer. Enzymes used for plasmid construction were purchased from NEB. Primers were obtained from IDT and Eurofins Genomics. gBlocks were obtained from IDT and Twist Bioscience. Gibson master mix was made according to the recipe at dx.doi.org/10.17504/protocols.io.n9xdh7n.

### Population genomic DNA sequencing

Aliquots (1 ml) of cultures archived during the evolution of JK1 had been maintained at -70 °C in M9/glucose containing 25% glycerol. Cells were harvested by scooping 400-600 µl of frozen culture into a 1.5 ml tube, thawing and centrifuging at 21,300 x g for 1 min at room temperature. Genomic DNA (gDNA) was extracted with an NEB Monarch gDNA kit (T3010) and submitted to SeqCenter for Illumina short-read sequencing. Reads were cleaned with Fastp 0.23.2^32^ using settings to trim the last base on each read and to remove reads with a phred quality less than 15 or with more than 20% unqualified bases, aligned with Bowtie2 2.1.0^33^ and analyzed with Breseq 0.35.4^34^ in polymorphism (mixed population) mode with CP009273.1 s the reference genome^35^. Read coverage exceeded 74x except for the final timepoint. For this sample, we sequenced gDNA prepared after growth of the culture overnight in LB to improve quantitation of *pykF* mutations, achieving 158x coverage. While the growth in LB might have resulted in some skewing of the population composition, the levels of the dominant mutations were comparable between sequence datasets (Supplementary Data Set 1).

The relative abundance of mutations in the evolving population of Δ*pdxB E. coli* that led to strain JK1 was determined by Breseq. (Frequencies of the *gapA\** and *ybhA\** mutations were determined in IGV^36^ when their abundances were too low to be registered by Breseq.) To simplify the analysis of the most important clones, we focused on mutations that were found at two or more consecutive timepoints, were present in over 10% of the reads or arose in a gene in which other mutations had already been found (Supplementary Data Set 1). Lineages were inferred based upon correlations between the relative abundances of mutations across multiple passages. The 206 kb amplification was identified based upon the read coverage map generated by Breseq (Fig. E1). The Muller diagram was generated in R 4.4.0 with the MullerPlot package^37^.

### Growth rate measurements

Cultures were routinely grown in M9/glucose (0.4% w/v) in the absence or presence of 10 µM pyridoxal. To measure growth rates in M9/glucose plus 10 µM pyridoxal, frozen glycerol stocks of *E. coli* strains were streaked onto LB agar plates and grown overnight at 37 °C. Single colonies were inoculated into 5 ml LB and grown overnight at 37 °C with shaking. Aliquots (250 µl) were inoculated into 25 ml of M9/glucose containing 10 µM pyridoxal in 40 mL vials in a FynchBio turbidostat. The cultures were grown at 37 °C with stirring (setting 8) and aeration provided by bubbling filtered air into the medium (25 ml/min). Cultures were started at an OD_600_ between 0.01 and 0.1 and then maintained at an OD_600_ between 0.14 and 0.16. (The FynchBio turbidostat measures IR light scattering rather than OD_600_. Standard curves were prepared to correlate light scattering to OD_600_.) The OD_600_ was recorded for at least 24 hours. Growth rates were determined using data from the turbidostat and a Python script (Related Manuscript File 1) that fits the OD_600_ for each dilution cycle (or each 1- to 2-hour interval if no dilution event occurred) to a single exponential. The µ values from multiple cycles (>6) were used to calculate the median for µ for each vial. Data reported are based on µ values for 7-13 biological replicates.

To measure growth rates in M9/glucose, starter cultures were grown in M9/glucose plus 10 µM pyridoxal as described above. After 18-24 h, cells were harvested by centrifugation at 4500 x g for 5 min at room temperature. Cell pellets were suspended in 1 ml of room-temperature M9/glucose. The suspensions were transferred to 1.5 ml tubes and centrifuged at 6000 x g for 1 min at room temperature to remove medium containing pyridoxal. The cells were washed 5 times with M9/glucose and suspended in 500 µl of M9/glucose before inoculation into 25 ml of M9/glucose in 40 ml vials in a FynchBio turbidostat to give an initial OD_600_ ≥ 0.14. The cultures were maintained in mid-log phase in the turbidostat as described above. OD_600_ was recorded for at least 72 hours. Growth rates were determined as described above. Slow-growing strains showed three distinct growth phases: 1) initial fast growth that we attribute to residual intracellular PLP; 2) a decline in OD_600_ that we attribute to lysis of some cells; and 3) consistent slow growth once the cells acclimate to conditions in which the only source of PLP is endogenous synthesis. We only included growth data from phase three when determining growth rates for these strains. Some biological replicates obtained mutations leading to faster growth; data from these replicates were excluded. Data reported are based on µ values for 3-12 biological replicates.

To measure growth rates in spent medium from the parental strain, 25 ml cultures of the parental Δ*pdxB* strain and the *gapA\** and *gapA* rpoS\** strains were started in 40 ml vials of a FynchBio turbidostat as described above. To result in enough spent medium and cells, four replicate cultures originating from the same colony of the *gapA\** strain and two of *gapA\** and *gapA* rpoS\** strains. After 24 h in M9/glucose, replicate cultures were combined and spent medium from the parental Δ*pdxB* strain was harvested by filtration through a 0.22 µm PES filter membrane. *gapA\** and *gapA* rpoS\** cells were harvested by centrifugating at 4500 x g for 5 min at room temperature. The pellets were suspended in 1 ml of room-temperature M9/glucose and transferred to a 1.5 ml tube. Half (500 µl) of the cell suspension was inoculated into 25 ml of spent medium and the other half was inoculated into 25 ml of M9/glucose. These cultures were grown in the turbidostat as described above using either spent medium or M9/glucose to dilute cultures when they reached an OD_600_=0.16. OD_600_ was recorded for at least 48 hours and growth rates were determined as described above. Data reported are based on µ values for 3-4 biological replicates.

### RNA sequencing

Aliquots (500 µl) of cells were withdrawn from the turbidostat after 48 h of growth in M9/glucose, added to 1 ml of bacterial protect reagent (Qiagen 76526), mixed by vortexing and incubated at room temperature according to the manufacturer’s protocol. The suspended cells were centrifuged for 10 min at 5000 x g at room temperature. The supernatants were discarded and the cell pellets were flash frozen and maintained at -70 °C. Subsequently, cell pellets were thawed at room temperature and RNA was extracted with a RNeasy Mini Kit (Qiagen 74104) according to the manufacturer’s protocol. Purified RNA samples were submitted to SeqCenter for Illumina short-read sequencing. Reads were cleaned with Fastp 0.23.2 using settings to remove reads with phred quality less than 15 or with more than 20% unqualified bases and aligned with Bowtie2 2.1.0. Bowtie alignment files were converted to sorted bam files with SAMtools^38^. Read counts for each gene were produced by HTSeq^39^. Data analysis and visualization were performed with edgeR^40^. Data are provided in Supplementary Data Set 2.

### Microscopy

Ten µl of 0.01% poly-L-lysine hydrobromide solution (PLL, MP Biomedicals 150176) was pipetted onto the center of a Premium Superfrost Plus glass slide (VWR 48311-601) and spread over a 1.5 x 1.5 cm square with a plastic cell spreader. Slides were dried at room temperature for one hour. One ml diH_2_O was used to wash the slide and remove excess PLL. Slides were dried and stored at room temperature for no more than three days.

Aliquots (200 µl) of cells were sampled from the turbidostat after a stable growth rate had been achieved and placed onto the middle of a prepared slide. Cells were immobilized on the surface for 20 minutes. Unbound cells were removed by washing the slide with 1 ml M9/glucose. M9/glucose (200 µl) containing 3.4 µM Syto9 and 40 µM propidium iodide (Invitrogen L7012) was pipetted onto the cell-containing region of the slide. The slides were allowed to sit for 5 min in a plastic light-proof box. The staining reagents were removed by washing the slide with 1 ml M9/glucose and the cell-containing region was covered by a VWR glass coverslip No 1.5 (48366-227). Prepared slides were maintained in the dark at room temperature and imaged within 1 h on a Yokogawa/Olympus CV1000.

### Quantification of vitamin B_6_ in cell lysates and spent media

Total vitamin B_6_ content in cell lysates and spent media was determined by a modification of a literature procedure to enzymatically convert all B_6_ vitamers to 4-pyridoxolactone (4-PLA, Fig. S1), a highly fluorescent molecule, prior to separation and quantitation by HPLC^20^. Cells were washed and grown as described above for determination of growth rates. Samples were harvested from the turbidostat after stable growth rates were achieved in M9/glucose. Three aliquots (7.5 ml) were withdrawn from each vial with sterile 10 ml syringes. Three smaller aliquots (100 µl) were also withdrawn and plated for enumeration of colony-forming units (CFUs) (described below). The samples were transferred to a cool (15 °C) dimly lit room for processing to minimize photodecomposition of B_6_ vitamers^41,42^. Cultures were pushed through a 0.22 µm MCE 25 mm syringe filter (CellTreat CT-229750) and ∼1 ml of spent medium was collected in an opaque black tube. Cells were then collected by washing the filter in the reverse direction using 1 ml 10 mM CHES buffer, pH 9.0. The cell suspension was transferred to a tube containing 1200 mg of 100 µm zirconium beads (Ops Diagnostics 10010026). An aliquot (50 µl) was removed and diluted in 450 µl of M9/glucose to quantify OD_600_ and CFUs. The spent media and cell suspensions were stored at -70 °C until processing for quantification of B_6_ vitamers.

Frozen cell suspensions with 1200 mg of 100 µm zirconium beads (Ops Diagnostics 10010026) were thawed in a 15 °C water bath for 5 min. Cells were lysed by shaking at full speed for 20 min on a horizontal microtube holder attachment (SI-H524) mounted on a Vortexer 2.0. Lysates were centrifuged for 1 min at 6000 x g and the supernatants were filtered through a 0.22 µm MCE 13 mm syringe filter (CellTreat CT-229750). An aliquot (200 µl) of the filtered cell lysate was placed in an opaque black tube containing 20 µl of premixed solution comprised of His-tagged recombinant Proteinase K (Abcam ab281339), 2.5 U His-tagged Benz-Neburase (GenScript Z03626-100), 5.5 mM MgCl_2_ and 550 mM ammonium acetate, pH 9.0. The reaction mixture was incubated in a water bath for 2 h at 37 °C to digest proteins and release covalently attached PLP from lysine residues. The remaining lysate was used for DNA and protein quantitation (described below). After protease digestion, 5 µl of 45 mM phenylmethylsulphonyl fluoride (PMSF) dissolved in ethanol was added and samples were tumbled for 10 min to inactivate proteases. An aliquot (165 µl) of 50 mM ammonium acetate, pH 9.0, containing 2 mM pyruvate, 2 mM oxidized nicotinamide adenine dinucleotide (NAD^+^), 5 µM flavin adenine dinucleotide (FAD), 0.5 mM MgCl_2_, 50 U recombinant shrimp alkaline phosphatase (rSAP, NEB M0371L), 0.14 µM pyridoxine oxidase (PNOX) and 0.22 µM pyridoxamine-pyruvate aminotransferase (PPAT) was added to the samples, followed by 60 µl of 0.53 µM PLDH. The resulting mixture was incubated in a 37 °C water bath for 1 h to convert B_6_ vitamers to 4-PLA. Purification and assays of PLDH, PPAT and PNOX are described below. His-tagged proteinase K (20 µl of 10 mg/ml) was added and the reaction mixture was incubated in a 37 °C water bath for 1 h to digest the conversion enzymes PNOX, PPAT and PLDH. Ni-NTA resin (10 µl, Cube Biotech 74105) was added to bind the His-tagged proteinase K and the reaction mixture was tumbled for 10 min. Finally, the reaction was filtered through a 0.22 µm MCE 13 mm syringe filter (CellTreat CT-229750) into 400 µl glass inserts (Agilent 5181-3377), placed in HPLC vials and capped with a septum cap.

Spent media samples were prepared and B_6_ vitamers were converted to 4-PLA as follows. Frozen spent media samples were thawed in a 15 °C water bath for 5 min and an aliquot (195 µl) was placed in an opaque black tube. EDTA (22 µl of 100 mM, pH 9.0) was mixed with the spent medium followed by 8 µl of 1 M NaOH. An aliquot (165 µl) of 100 mM ammonium acetate, pH 9.0, containing 2 mM pyruvate, 2 mM oxidized nicotinamide adenine dinucleotide (NAD^+^), 5 µM flavin adenine dinucleotide (FAD), 1 mM MgCl_2_, 50 U recombinant shrimp alkaline phosphatase (rSAP, NEB M0371L), 0.14 µM pyridoxine oxidase (PNOX) and 0.22 µM pyridoxamine-pyruvate aminotransferase (PPAT) was added to the tumbled samples, followed by 60 µl of 0.53 µM PLDH. The reaction was incubated in a 37 °C water bath for 2 h to convert B_6_ vitamers to 4-PLA. Conversion enzymes were digested with His-tagged proteinase K and the solution was filtered as described above.

4-PLA was quantified by HPLC. The HPLC system consisted of an Agilent 1100 degasser (G1379A), a quat pump (G1311A), an autosampler (G1367A) with a 1290 thermostat (G1330B) and a fluorescence detector (G1321A). An Infinitylab Poroshell 120EC-C18 column (4.6 x 250 mm, 2.7 µm, Agilent 690975-902) with an Infinitylab Poroshell 120EC-C18 guard column (4.6 x 5 mm, 2.7 µm, Agilent 820750-911) was used for separation of compounds. All samples were maintained in the dark at 4 °C in the autosampler prior to injection on the column. A standard curve of 4-PLA (Sigma 02582) from 0 to 40 nM was run for each set of 16 samples. After injection of 40 µl samples, 4-PLA was eluted with 2 column volumes of mobile phase A (10 mM ammonium acetate, pH 8.75, containing 5% (v/v) acetonitrile at a flow rate of 0.75 ml/min. The column was washed with 5 column volumes of mobile phase B (95% (v/v) acetonitrile/water) and re-equilibrated with 3 column volumes of mobile phase A. The fluorescence intensity of the eluate was monitored at 425 nm (excitation at 355 nm) with a gain of 13 and response time of 4 s. Peaks were automatically integrated by Chemstation.

### Enumeration of colony forming units (CFUs)

Aliquots (100 µl) of diluted cultures were spread on LB agar plates and incubated at 37 °C for 12-16 h. Plates were imaged with a digital camera in an imaging box^43^ and colonies were counted with a custom Matlab script (Related Manuscript File 2). Plates with fewer than 10 colonies and more than 2000 colonies were excluded from CFU calculations. OD_600_ and CFU were linearly correlated (3.52 x 10^8^ CFU/OD_600_, r^2^ = 0.76) and did not change between cell types, allowing the number of CFUs to be calculated from OD_600_.

### Quantification of DNA in lysates and spent medium

DNA in lysate and spent medium samples was quantified using the Qubit HS DNA Detection Kit (ThermoFisher Q33231) in a 96-well plate. A solution comprised of 0.5 µl Component A and 89.5 µl Component B was mixed with an aliquot (10 µl) of the sample or DNA standard (concentrations ranging from 0.25 to 6 µmol/ml for lysate samples or 0.0078 to 6 µmol/ml for spent medium samples) and emission at 532 nm (excitation at 502 nm) was measured in a Thermo Varioscan 3.01.15 plate reader. The fluorescence of the standards was fitted to a quadratic curve and the regression coefficients were used to calculate the DNA concentrations in the samples.

### Quantification of protein in lysates

Protein in lysates was quantified using a Coomassie protein assay in a 96-well plate. An aliquot (75 µl) of Coomassie protein assay reagent (Thermo 1856209) was mixed with 25 µl of the sample or BSA standard (BioRad 500-0207) (concentrations ranging from 2.5 to 40 µmol/ml) and absorbance at 595 nm was measured in a Thermo Varioscan 3.01.15 plate reader. The standards were fitted to a quadratic curve and regression coefficients were used to calculate the protein concentrations in the samples.

### Protein purification

gBlocks encoding enzymes required for analysis of B_6_ vitamers were ordered from Twist Bioscience. The gBlock encoding PLDH was assembled into a pET28 vector with a His_10_-SUMO tag incorporated at the N-terminus (pKAW077, Table S2). The gBlocks encoding PNOX and PPAT were cloned into a pET46 vector with a His_6_ tag incorporated at the N-terminus (pKAW110 and pKAW111, Table S2). The expression plasmids were introduced into *E. coli* NiCo21(DE3) (Table S1) by electroporation.

Expression and purification of PLDH was carried out as follows. A pipette tip was used to inoculate a small quantity of a frozen culture of *E. coli* NiCo21(DE3) containing pKAW077 into 10 mL of LB containing 50 µg/mL kanamycin in a 50 mL flask. The cultures were grown at 37 °C overnight with shaking and then inoculated into one L of TB containing 50 µg/mL kanamycin. After reaching an OD_600_ of 0.5-0.6, the culture was cooled to room temperature and isopropyl β-D-1-thiogalactopyranoside (IPTG) was added to a final concentration of 1 mM induce protein expression. The culture was maintained at 25 °C with shaking for 16 h and cells were harvested by centrifugation at 6000 x g for 20 min at 4 °C. The cell pellet was suspended in 35 ml cold purification buffer with protease inhibitor cocktail (Thermo Scientific™ A32955) and lysozyme (1 mg/ml) and incubated for 1 h at 4 °C with tumbling. PLDH purification buffer consisted of 20 mM potassium phosphate buffer, pH 8.0, containing 0.1% (v/v) 2-mercaptoethanol, 10% (v/v) glycerol and 1 mM EDTA^44^. Cells were lysed by sonication (5 cycles, 10 s on, 20 s off) on ice. The disrupted cells were centrifuged at 20,000 x g at 4 °C for 20 min to remove cell debris. Protein was purified using a hand-packed Ni-NTA resin column (300 µl bed volume of Cube Biotech 74105) using a step gradient of PLDH purification buffer containing increasing amounts of imidazole (10 ml of 20 mM, 5 ml of 50 mM, 5 ml of 100 mM, 5 ml of 200 mM and 5 ml of 500 mM). SDS-PAGE was used to identify fractions containing PLDH. Fractions from the 200 mM and 500 mM imidazole elutions were combined. PLDH was further purified by size exclusion chromatography (Superdex 200 10/300 GL, 24 ml bed volume) with PLDH storage buffer (20 mM sodium carbonate, pH 9.0, containing 0.1% (v/v) 2-mercaptoethanol, 10% (w/v) glycerol and 1 mM EDTA) as a running buffer. The final protein concentration was calculated using the A_280_ and the PLDH extinction coefficient calculated with Expasy ProtParam (https://web.expasy.org/protparam/). Purified protein was flash frozen and stored at -70 °C.

Expression of PPAT and harvesting of cells was carried out as described above for PLDH. PPAT purification buffer A consisted of 20 mM sodium phosphate, pH 7.4, containing 300 mM NaCl. Cells were lysed by sonication (5 cycles, 10 s on, 20 s off) on ice and centrifuged at 20,000 x g at 4 °C for 20 min to remove cell debris. Residual cell debris was removed by filtration through a 0.45 µm 33 mm PVDF syringe filter (Millipore SLHVR33RS). Proteins were purified with a 1 ml HisTrap FF (Cytiva 17531901) column on an AKTA FLPC using a 30 ml linear gradient of PPAT purification buffer A and buffer B (buffer A containing 500 mM imidazole). Protein elution was monitored by OD_280_ and fractions containing PPAT were identified by SDS-PAGE. PPAT was further purified by size exclusion chromatography (Superdex 200 10/300 GL, 24 ml bed volume) with PPAT purification buffer A (20 mM sodium phosphate, pH 7.4, containing 300 mM NaCl) as a running buffer. Protein elution was monitored by A_280_ and fractions containing PPAT were identified by SDS-PAGE. The final protein concentration was calculated using the A_280_ and the extinction coefficient calculated with Expasy ProtParam. Purified protein was flash frozen and stored at -70 °C.

Expression of PNOX required the chaperones GroEL and GroES^45^, which were overexpressed from pGro7 (Takara Bio 3340). Purification of PNOX proceeded as described above with a few modifications. Overnight starter cultures of *E. coli* NiCo21(DE3) containing pKAW111 and pGro7 were grown in LB containing 50 µg/ml kanamycin to maintain the overexpression plasmid and 30 µg/ml chloramphenicol to maintain pGro7; expression cultures also contained 50 µg/ml kanamycin and 30 µg/ml chloramphenicol. The cell pellet was suspended in PNOX purification buffer (20 mM potassium phosphate buffer, pH 8.0, containing 300 mM NaCl, 5 µM FAD, 0.1% (v/v) 2-mercaptoethanol, 10% (w/v) glycerol, 1 mM EDTA and 0.01% (v/v) Tween 20, as described previously^45^). Cells were lysed and separated from cell debris as described above. Protein was purified using a hand-packed benchtop Ni-NTA resin column (300 µl bed volume of Cube Biotech 74105) using a step gradient of PLDH purification buffer containing increasing amounts of imidazole (10 ml of 20 mM, 5 ml of 50 mM, 5 ml of 100 mM, 5 ml of 200 mM and 5 ml of 500 mM). SDS-PAGE was used to identify fractions containing PLDH. Fractions from the 100 mM and 200 mM imidazole elutions were combined. Buffer exchange was performed by several steps of concentration by centrifugation at 6000 x g for 20 min at 4 °C with a 30 kDa-cutoff membrane centrifugal filter (Millipore UFC9030) and dilution in PNOX storage buffer (50 mM Tris buffer, pH 8.0, containing 5 µM FAD, 0.1% (v/v) 2-mercaptoethanol, 10% (w/v) glycerol, 1 mM EDTA and 0.01% (v/v) Tween 20). Purified protein was flash frozen and maintained at -70 °C.

### Enzyme assays

The activities of PLDH, PPAT, PNOX and rSAP were measured under the conditions used to convert B_6_ vitamers to 4-PLA (50 mM ammonium acetate, pH 9.0, containing 0.5 mM MgCl_2_, 2.5 µM FAD, 1 mM NAD^+^ and 1 mM pyruvate). Reaction mixtures were incubated in wells of a 96-well flat-bottom plate in a Varioskan plate reader for 5 min while measuring the background change in OD_340_ every min. Reactions were initiated by addition of PL, PM, PN or PLP to a final concentration of 1 mM. This assay procedure was used to determine the amount of each enzyme required to convert >99% of the B_6_ vitamers to 4-PLA in 30 min. All enzymes were stable at -80 °C for at least 6 months.

### Genome editing

A helper plasmid (pDY118A) encoding Cas9, I-SceI and the λ Red genes (alpha, beta and gam) was transformed into Δ*pdxB*::*kanR E. coli* BW25113 from the Keio collection^46^ as described previously^47^. The four singly edited strains were constructed from the Δ*pdxB*::*kanR* strain as described below. Strains with multiple edits were constructed by successively adding edits to the singly edited strains.

We generated donor and guide plasmids containing I-SceI recognition sites and a *sacB* counter-selection marker from previously constructed donor plasmids pDonor1 and pDonor2^47^, respectively. Donor plasmids contain both an editing cassette and an sgRNA sequence, while guide plasmids contain only an sgRNA sequence. The I-SceI recognition sites and *sacB* counter-selection marker were included to aid plasmid curing after genome editing. However, leaky expression of I-SceI from the pDY118A helper plasmid proved to be efficient enough to cure the donor and guide plasmids so that neither I-SceI induction nor SacB counter-selection was necessary. PCR amplicons, gBlocks and annealed primers used for Gibson assembly of donor and guide plasmids for genome editing are listed in Table S7. All PCRs were carried out using the manufacturer’s protocol for NEB Q5 High-Fidelity DNA polymerase M0492L. Assembled plasmids were transformed into DH5a by electroporation. Correct plasmid construction was verified by whole plasmid sequencing before genome editing procedures.

Cas9-assisted lambda Red recombination was used to introduce the 3812 bp deletion (Δ). A guide plasmid encoding an sgRNA targeting *ybhA* was assembled from the backbone of pDonor1-SacB, excluding the editing cassette, and a sgRNA sequence made by annealing two complimentary 60-nt primers (Tables S4 and S7), resulting in a 20 bp ds sgRNA sequence flanked by 20 bp extensions for Gibson assembly. Editing was performed as described previously^47^ except that 30 µg/ml chloramphenicol was used to maintain pDY118A rather than 34 µg/ml. Editing was confirmed by colony PCR and Sanger sequencing.

Two-step Cas9-assisted lambda Red recombination was used to introduce the *gapA\** and *rpoC\** mutations. Because these genes are essential, we introduced multiple synonymous mutations into the genes in the first step of editing to maintain the essential function. In the second step, the region containing the multiple synonymous mutations was targeted by an sgRNA to enable replacement with the edited sequence provided as a linear editing cassette (rather than as a cassette in a donor plasmid). Round 1 donor plasmids used to introduce multiple synonymous mutations into the genome within 20 bp of the intended edits were constructed by Gibson assembly of the backbone of pDonor2-SacB with a gBlock (IDT) containing the synonymously recoded gene segment and extensions required for Gibson assembly (Table S7). In the second step, a guide plasmid was used to target the synonymously recoded site in the genome. Guide plasmids were constructed by Gibson assembly of a backbone amplified from pDonor1-SacB (excluding the previous sgRNA sequence) with an sgRNA sequence made by annealing two complimentary oligonucleotides (Table S4). Linear editing cassettes were amplified from JK1 gDNA using primers KW019 and KW020 for *gapA\** and KW013 and KW014 for *rpoC\** (Table S3). Editing was performed as described previously^47^ except that 100 µg/ml streptomycin was used for selection of guide plasmids rather than 100 µg/ml ampicillin.

I-SceI-assisted editing^48^ was used to introduce the *rpoS\** mutation. Edits were introduced using a three-part dsDNA editing cassette comprised of: first, a segment containing a homology arm (HA) upstream of the edit, the edit and a short downstream HA; second, a segment containing an 18-bp I-SceI cut site and a streptomycin resistance gene; and third, a final segment containing a short HA upstream of the edit, the edit and a downstream HA. The editing cassette was amplified from pKAW066 using the manufacturer’s protocol for NEB Q5 Hot Start DNA Polymerase (M0494L) and primers KW059 and KW385 (Table S3). Lambda Red proteins were expressed and cells were prepared for electroporation as described previously. The editing cassette (100 ng) was introduced into washed competent cells (50 µl) by electroporation, after which the cells were added to 1 ml SOC at 30 °C and allowed to recover for 3 h and then plated on LB agar plates containing 30 μg/ml chloramphenicol to maintain pDY118A and 50 μg/ml streptomycin to select for cells that had successfully recombined the editing cassette into the genome. After overnight incubation at 30 °C, colonies were streaked onto LB agar plates containing 30 μg/ml chloramphenicol and 50 μg/ml streptomycin and grown overnight at 30 °C to ensure elimination of unedited cells. Individual colonies were then streaked onto fresh LB agar plates containing 30 μg/ml chloramphenicol and 1 mM IPTG to induce I-SceI expression and promote RecA-mediated recombination to obtain the scarless edit. After overnight incubation at 30 °C, colonies were screened for incorporation of the desired edit.

Plasmids were cured from edited colonies by overnight growth in 5 ml LB at 37 °C with shaking, followed by streaking onto LB agar plates. After overnight growth on LB agar plates at 37 °C, individual colonies were streaked onto LB plates containing: 1) no antibiotics; 2) 30 µg/ml chloramphenicol to test for retention of the helper plasmid pDY118A; and 3) 100 µg/ml ampicillin or streptomycin to test for retention of donor and guide plasmids, respectively. Plates were incubated at 37 °C overnight. Colonies that grew on LB agar, but not on LB agar containing antibiotics, were picked into 5 ml LB and grown overnight at 37 °C with shaking. An aliquot of the culture (750 µl) was diluted with 750 µl of sterile 50% glycerol, frozen and maintained at -70 °C. Cells were harvested from the remaining culture by centrifugation at 4500 x g for 5 min at room temperature and used for gDNA purification as described above. Whole-genome sequencing was carried out in parental Δ*pdxB*::*kanR E. coli* BW25113 and the edited strains at SeqCenter and SeqCoast to ensure correct editing and the absence of unintended mutations elsewhere in the genome.

## Supporting information

Related Manuscript File 1 - Python script

Related Manuscript File 2 - Matlab script

Supplementary Data Set 1 - Mutations in evolving population

Supplementary Data Set 2 - RNA Seq data

## Extended Data

**Fig. E1.**
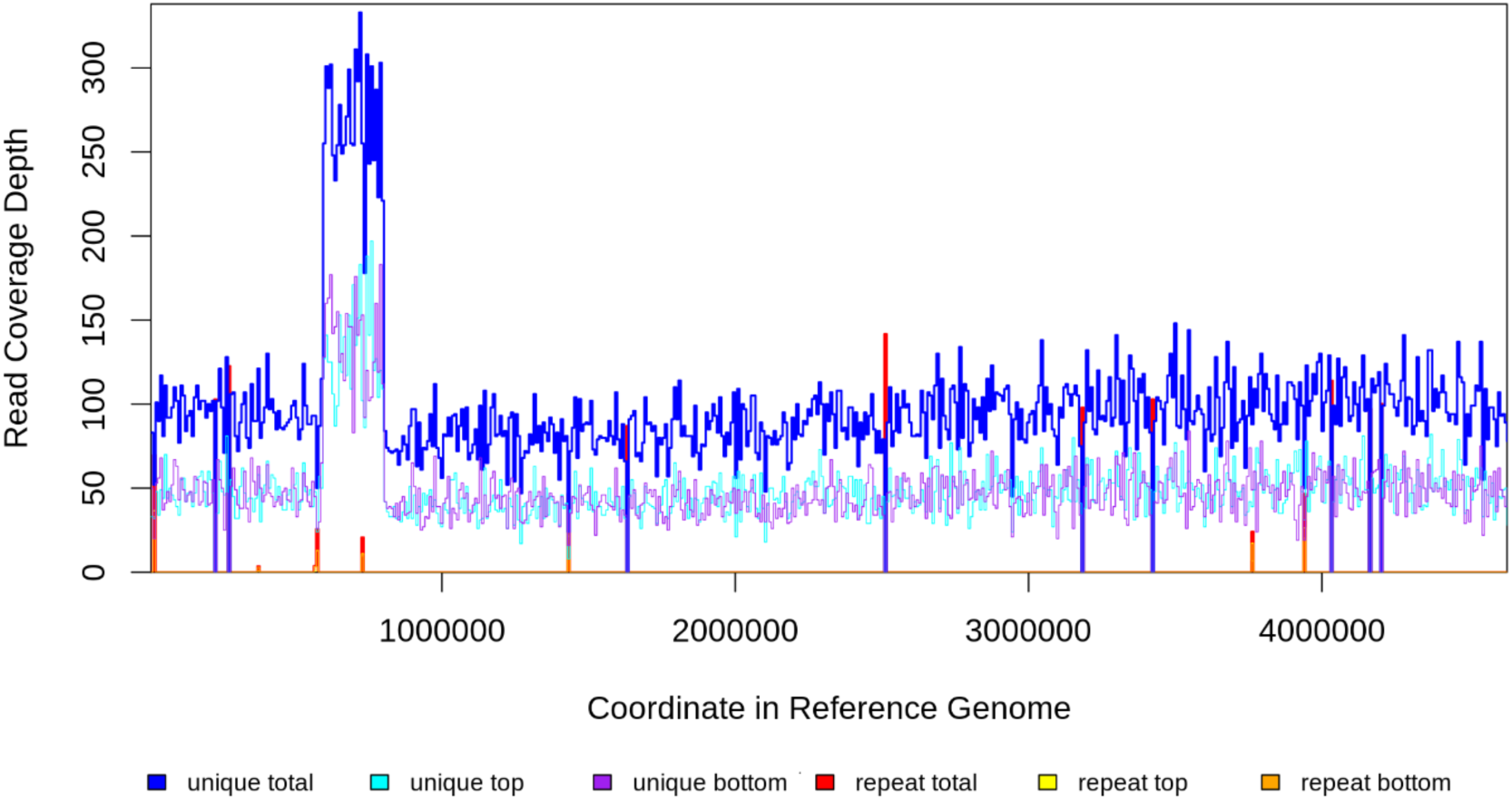
Read coverage in population gDNA at passage 8 when the clone containing the amplification dominated the population. Read coverage averages 92 but is elevated to ∼275 between genome coordinates 592434 and 798903.

**Fig. E2.**
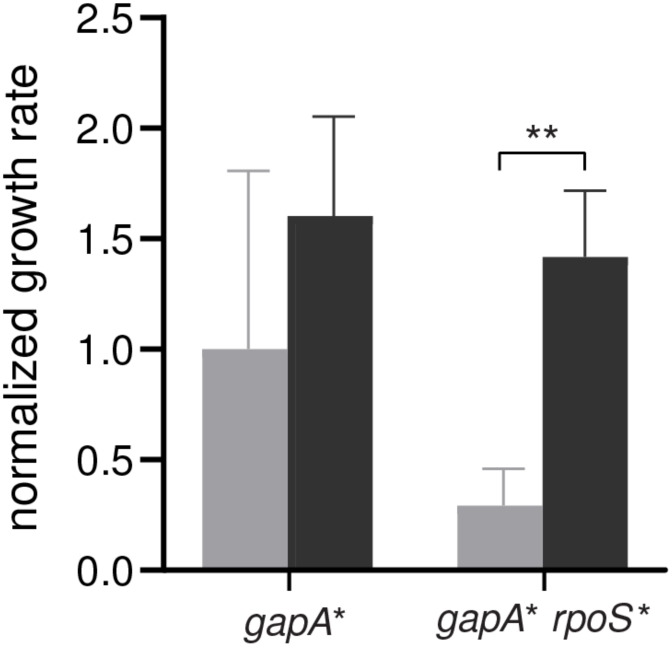
Spent medium collected after growth of Δ*pdxB E. coli* improves growth of the *gapA* rpoS\** strain. Light grey, growth rate in M9/glucose; dark grey, growth rate in medium collected after growth of Δ*pdxB E. coli* to OD_600_ = 0.15 in M9/glucose. Error bars represent 1 standard deviation. P-values adjusted by Dunnett T3 correction for multiple comparisons (Graphpad Prism 8). *\*\*,* p-adj < 0.01.

**Fig. E3.**
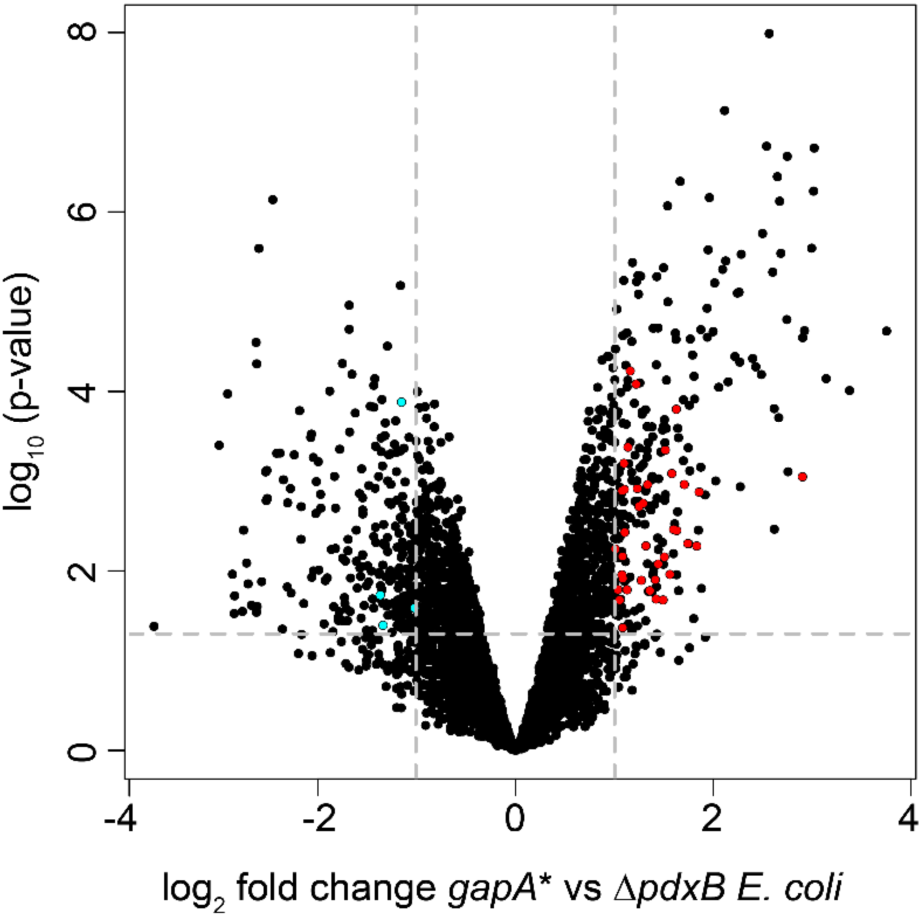
The *gapA\** mutation in the Δ*pdxB* background causes significant changes in gene expression. Dark red, genes in the RpoS regulon that are significantly upregulated; cyan, genes in the RpoS regulon that are significantly downregulated. Horizontal grey dashed line, cutoff for statistical significance with p-adj < 0.05. Vertical grey dashed lines, cutoff for changes in expression greater than 2-fold.

## Acknowledgements

This work was supported by NIH R01GM135364 to SDC and a Cooperative Research in Environmental Sciences graduate student fellowship to KAW.

## Peer review

This work has not yet been peer-reviewed.

## References

1. Koonin, E. Comparative genomics, minimal gene sets, and the last universal common ancestor. Nature Rev. Microbiol. 1, 127–136 (2003).

2. Moody, E.R.R. et al. The nature of the last universal common ancestor and its impact on the early Earth system. Nat Ecol Evol (2024).

3. de Souza, M.L., Seffernick, J., Martinez, B., Sadowsky, M.J. & Wackett, L.P. The atrazine catabolism genes *atzABC* are widespread and highly conserved. J. Bacteriol. 180, 1951–1954 (1998).

4. Copley, S.D. et al. The whole genome sequence of *Sphingobium chlorophenolicum* L-1: insights into the evolution of the pentachlorophenol degradation pathway. Genome Biol Evol 4, 184–98 (2012).

5. Poelarends, G.J., Wilkens, M., Larkin, M.J., van Elsas, J.D. & Janssen, D.B. Degradation of 1,3-dichloropropene by *Pseudomonas cichorii* 170. Appl Environ Microbiol 64, 2931–6 (1998).

6. Dong, Z. et al. Mechanism for biodegradation of sulfamethazine by *Bacillus cereus* H38. Sci Total Environ 809, 152237 (2022).

7. Ferguson, J.F. & Pietari, J.M. Anaerobic transformations and bioremediation of chlorinated solvents. Environ Pollut 107, 209–15 (2000).

8. Das, S., Cherwoo, L. & Singh, R. Decoding dye degradation: Microbial remediation of textile industry effluents. Biotechnol Notes 4, 64–76 (2023).

9. Esteve-Nunez, A., Caballero, A. & Ramos, J.L. Biological degradation of 2,4,6-trinitrotoluene. Microbiol Mol Biol Rev 65, 335–52, table of contents (2001).

10. Teichmann, S.A. et al. The evolution and structural anatomy of the small molecule metabolic pathways in *Escherichia coli*. J Mol Biol 311, 693–708 (2001).

11. Gerlt, J.A., Babbitt, P.C. & Rayment, I. Divergent evolution in the enolase superfamily: the interplay of mechanism and specificity. Arch. Biochem. Biophys. 433, 59–70 (2005).

12. Burroughs, A.M., Allen, K.N., Dunaway-Mariano, D. & Aravind, L. Evolutionary genomics of the HAD superfamily: understanding the structural adaptations and catalytic diversity in a superfamily of phosphoesterases and allied enzymes. J Mol Biol 361, 1003–34 (2006).

13. Roodveldt, C. & Tawfik, D.S. Shared promiscuous activities and evolutionary features in various members of the amidohydrolase superfamily. Biochemistry 44, 12728–36 (2005).

14. Bergthorsson, U., Andersson, D.I. & Roth, J.R. Ohno’s dilemma: evolution of new genes under continuous selection. Proc. Natl. Acad. Sci. U S A 104, 17004–9 (2007).

15. D’Ari, R. & Casadesus, J. Underground metabolism. Bioessays 20, 181–6 (1998).

16. Kim, J. et al. Hidden resources in the *Escherichia coli* genome restore PLP synthesis and robust growth after deletion of the essential gene pdxB. Proc Natl Acad Sci U S A 116, 24164–24173 (2019).

17. Cheng, K.K. et al. Global metabolic network reorganization by adaptive mutations allows fast growth of *Escherichia coli* on glycerol. Nat Commun 5, 3233 (2014).

18. Conrad, T.M. et al. RNA polymerase mutants found through adaptive evolution reprogram *Escherichia coli* for optimal growth in minimal media. Proc Natl Acad Sci U S A 107, 20500–5 (2010).

19. Sandberg, T.E. et al. Evolution of *Escherichia coli* to 42 degrees C and subsequent genetic engineering reveals adaptive mechanisms and novel mutations. Mol Biol Evol 31, 2647–62 (2014).

20. Thi Viet Do, H., Ide, Y., Mugo, A.N. & Yagi, T. All-enzymatic HPLC method for determination of individual and total contents of vitamin B(6) in foods. Food Nutr Res 56(2012).

21. Morgenthaler, A.B. et al. Mutations that improve efficiency of a weak-link enzyme are rare compared to adaptive mutations elsewhere in the genome. Elife 8(2019).

22. Payen, C. et al. High-throughput identification of adaptive mutations in experimentally evolved yeast populations. PLoS Genet 12, e1006339 (2016).

23. Ferenci, T. The spread of a beneficial mutation in experimental bacterial populations: the influence of the environment and genotype on the fixation of *rpoS* mutations. Heredity (Edinb) 100, 446–52 (2008).

24. Knoppel, A. et al. Genetic adaptation to growth under laboratory conditions in *Escherichia coli* and *Salmonella enterica*. Front Microbiol 9, 756 (2018).

25. Maharjan, R., Seeto, S., Notley-McRobb, L. & Ferenci, T. Clonal adaptive radiation in a constant environment. Science 313, 514–7 (2006).

26. Notley-McRobb, L., King, T. & Ferenci, T. *rpoS* mutations and loss of general stress resistance in *Escherichia coli* populations as a consequence of conflict between competing stress responses. J Bacteriol 184, 806–11 (2002).

27. Robinson, T., Smith, P., Alberts, E.R., Colussi-Pelaez, M. & Schuster, M. Cooperation and cheating through a secreted aminopeptidase in the *Pseudomonas aeruginosa* RpoS response. mBio 11(2020).

28. Sugimoto, R., Saito, N., Shimada, T. & Tanaka, K. Identification of YbhA as the pyridoxal 5’-phosphate (PLP) phosphatase in *Escherichia coli*: Importance of PLP homeostasis on the bacterial growth. J Gen Appl Microbiol 63, 362–368 (2018).

29. Wytock, T.P. et al. Experimental evolution of diverse *Escherichia coli* metabolic mutants identifies genetic loci for convergent adaptation of growth rate. PLoS Genet 14, e1007284 (2018).

30. Bouillet, S., Bauer, T.S. & Gottesman, S. RpoS and the bacterial general stress response. Microbiol Mol Biol Rev 88, e0015122 (2024).

31. Woods, R., Schneider, D., Winkworth, C.L., Riley, M.A. & Lenski, R.E. Tests of parallel molecular evolution in a long-term experiment with *Escherichia coli*. Proc Natl Acad Sci U S A 103, 9107–12 (2006).

32. Chen, S., Zhou, Y., Chen, Y. & Gu, J. fastp: an ultra-fast all-in-one FASTQ preprocessor. Bioinformatics 34, i884–i890 (2018).

33. Langmead, B. & Salzberg, S.L. Fast gapped-read alignment with Bowtie 2. Nat Methods 9, 357–9 (2012).

34. Deatherage, D.E. & Barrick, J.E. Identification of mutations in laboratory-evolved microbes from next-generation sequencing data using breseq. Methods Mol Biol 1151, 165–88 (2014).

35. Grenier, F., Matteau, D., Baby, V. & Rodrigue, S. Complete genome sequence of *Escherichia coli* BW25113. Genome Announc 2(2014).

36. Thorvaldsdottir, H., Robinson, J.T. & Mesirov, J.P. Integrative Genomics Viewer (IGV): high-performance genomics data visualization and exploration. Brief Bioinform 14, 178–92 (2013).

37. Farahour, F., Saeedghalati, M. & Hoffman, D. MullerPlot: Generates Muller plot from population/abundance/frequency dynamics data. 0.1.3 edn R package (2022).

38. Danecek, P. et al. Twelve years of SAMtools and BCFtools. Gigascience 10(2021).

39. Putri, G.H., Anders, S., Pyl, P.T., Pimanda, J.E. & Zanini, F. Analysing high-throughput sequencing data in Python with HTSeq 2.0. Bioinformatics 38, 2943–2945 (2022).

40. Robinson, M.D., McCarthy, D.J. & Smyth, G.K. edgeR: a Bioconductor package for differential expression analysis of digital gene expression data. Bioinformatics 26, 139–40 (2010).

41. Saidi, B. & Warthesen, J.J. Influence of pH and light on the kinetics of vitamin B6 degradation. Agricultural Food Chem 31, 878–880 (1983).

42. Reiber, H. Photochemical reactions of vitamin B_6_ compounds, isolation and properties of products. Biochim Biophys Acta 279, 310–5 (1972).

43. Smith, P. & Schuster, M. Inexpensive apparatus for high-quality imaging of microbial growth on agar llates. Front Microbiol 12, 689476 (2021).

44. Yokochi, N., Nishimura, S., Yoshikane, Y., Ohnishi, K. & Yagi, T. Identification of a new tetrameric pyridoxal 4-dehydrogenase as the second enzyme in the degradation pathway for pyridoxine in a nitrogen-fixing symbiotic bacterium, *Mesorhizobium loti*. Arch Biochem Biophys 452, 1–8 (2006).

45. Yuan, B., Yoshikane, Y., Yokochi, N., Ohnishi, K. & Yagi, T. The nitrogen-fixing symbiotic bacterium *Mesorhizobium loti* has and expresses the gene encoding pyridoxine 4-oxidase involved in the degradation of vitamin B6. FEMS Microbiol Lett 234, 225–30 (2004).

46. Baba, T. et al. Construction of *Escherichi coli* K-12 in-frame, single-gene knock-out mutants -- the Keio collection. Mol. Systems Biol. 2, Article number 2006.0008 (2006).

47. Yang, D.D., Rusch, L.M., Widney, K.A., Morgenthaler, A.B. & Copley, S.D. Synonymous edits in the *Escherichia coli* genome have substantial and condition-dependent effects on fitness. Proc Natl Acad Sci U S A 121, e2316834121 (2024).

48. Kim, J., Webb, A.M., Kershner, J.P., Blaskowski, S. & Copley, S.D. A versatile and highly efficient method for scarless genome editing in *Escherichia coli* and *Salmonella enterica*. BMC Biotechnol 14, 84 (2014).

