## Supplementary material for "A cheater founds the winning lineages during evolution of a novel metabolic pathway": Related Manuscript File 1 - Python script

import csv
import numpy as np
import matplotlib.pyplot as plt
from math import log2 as log2
from numpy import log as ln
import unicodedata
import re
from scipy.optimize import curve_fit
import os
from sklearn.linear_model import LinearRegression
from statistics import mean
from copy import copy


#Makes it easier to make output files with the desired names.
def slugify(value, allow_unicode=False):
    """
    Taken from https://github.com/django/django/blob/master/django/utils/text.py
    Convert to ASCII if 'allow_unicode' is False. Convert spaces or repeated
    dashes to single dashes. Remove characters that aren't alphanumerics,
    underscores, or hyphens. Convert to lowercase. Also strip leading and
    trailing whitespace, dashes, and underscores.
    """
    value = str(value)
    if allow_unicode:
        value = unicodedata.normalize('NFKC', value)
    else:
        value = unicodedata.normalize('NFKD', value).encode('ascii', 'ignore').decode('ascii')
    value = re.sub(r'[^\w\s-]', '', value.lower())
    return re.sub(r'[-\s]+', '-', value).strip('-_')


#Imports a .csv file with either the numbers or the names of the samples.

def importCSV(z):
    
    holder = []
    with open(z, newline = '') as csvfile:
        reader = csv.reader(csvfile)
        for row in reader:
            for x in range(len(row)):
                holder.append(row[x])
            
    return(holder)


#Asks the user to input which vials they used and what the samples are in each vial.
#It returns the numbers of the first and last vials used and an array containing the sample names associated with each vial.
def getinfo_():
    vialnums = []
    
    numberfilecheck = input("Do you have a CSV file with the vial numbers you used in one line?: (yes/no): ")
    if numberfilecheck == 'yes':
        numberfilename = input("What is the name of the file? (ex. filename.csv): ")
        vialnums_temp = importCSV(numberfilename)
        for n in range(len(vialnums_temp)):
            vialnums.append(int(vialnums_temp[n]))
                               
    else:
        consecutive = input("Are the vials that you would like to run consecutive? (ie, 3,4,5 or 2,3) (yes/no): ")                       
        if consecutive == 'yes':
            firstvial = int(input("Enter vial number for the first vial used: "))
            lastvial = int(input("Enter vial number for the last vial used: "))

            for n in range(firstvial, lastvial+1):
                vialnums.append(n)

        else:
            vialnums_temp = input("Enter the vial numbers you'd like analyzed in order with commas and no spaces separating them: ")
            vialnums_temp = vialnums_temp.split(',')
            for n in range(len(vialnums_temp)):
                vialnums.append(int(vialnums_temp[n])) 
                               
                               
    namesfilecheck = input("Do you have a CSV file with the names of strains in the vials you used in one line? (yes/no): ")
    if namesfilecheck == 'yes':
        namesfilename = input("What is the name of the file? (ex. filename.csv): ")
        samplenames = importCSV(namesfilename)
        
    else:
        samplenames = input("Enter the sample names in each tube in order with commas and no spaces separating them: ")
        samplenames = samplenames.split(',') 
    
    return(vialnums, samplenames)


#Imports the .csv OD file that is made by the turbidostat (includes ODs and the time at which the OD was measured).
#It takes in the vial number that is currently being analyzed
#It outputs an array of arrays containing times and ODs
#It skips the first row in the file since it contains headers
def importOD(z):
    
    vialOD = []
    with open('vial'+str(z)+'_OD.txt', newline = '') as csvfile:
        reader = csv.reader(csvfile)
        next(reader)
        for row in reader:
            vialOD.append([float(row[0]), float(row[1])])
            
    return(vialOD)


#Imports the .csv pump log that is made by the turbidostat (includes the time at which dilutions (pump events) occured).
#It takes in the vial number that is currently being analyzed
#It outputs an array of arrays containing times and ODs
#It skips the first row in the file since it contains headers
#It skips the second row since it is always [0,0]
def importpumptimes(z):
    
    vialpumptimes = []
    with open('vial'+str(z)+'_pump_log.txt', newline = '') as csvfile:
        reader = csv.reader(csvfile)
        next(reader)
        next(reader)
        for row in reader:
            vialpumptimes.append(float(row[0]))
        
    return(vialpumptimes)


#Applies a linear fit to the data contained in , with pumpgrowth[0][:] being the x values and pumpgrowth[1][:] being the y values
#Has to put the data contained in pumpgrowth into numpy array to do fit. In theory could instead just have pumpgrowth be a numpy array.
#Also graphs the fit onto ax1
#Returns the best fit slope coeficient (m in y=mx+b) and ax1 with the fit graphed on it
def func(x, a, b, c):
    return ((a * np.exp(b * x)) + c)


def dofit(pumpgrowth):

    xarray = np.array([])
    yarray = np.array([])
    xarray_orig = np.array([])
    
    xinit = pumpgrowth[0][0]
    yinit_med = np.median(np.array([pumpgrowth[0][1], pumpgrowth[1][1], pumpgrowth[2][1]]))
    xfinal = pumpgrowth[-1][0]
    yfinal_med = np.median(np.array([pumpgrowth[-1][1], pumpgrowth[-2][1], pumpgrowth[-3][1]]))
    
    for x in range(len(pumpgrowth)):
        xarray = np.append(xarray, pumpgrowth[x][0]-xinit)
        xarray_orig = np.append(xarray_orig, pumpgrowth[x][0])
        yarray = np.append(yarray, pumpgrowth[x][1])
        
    growthrateguess = abs( ( ln(yfinal_med)-ln(yinit_med) ) / (xfinal - xinit) )


    popt, pcov = curve_fit(func, xarray, yarray, maxfev = 10000, p0 = (yinit_med,growthrateguess,0), bounds = ([yinit_med - 0.01, 0, -0.01], [yinit_med + 0.01,growthrateguess + 0.8,0.01]))

    #ax1.scatter(xarray_orig, yarray)
    #ax1.plot(xarray_orig, func(xarray, *popt),color='k')

    
    
    return(popt[1])
    
    
#Calculates the growth rate for each section between pumping events
#Takes in the vialOD and vialpumptimes array that were made by importing the two CSVs
#Takes in ax1 that was made in the main code
#Requires dofit() that was defined above
#Returns an array "times" that has the times at which or immediately after a pump event occurs
#Returns an array "onlypumpgrowthrate" that has the growth rates from each pump event
#"times" and "onlypumpgrowthrate" are of equal size, and the time can be associated with the growth rate of the section immediately before that pump event
#Could instead have made these a numpy array
#Also returns ax1, now with the ln(OD600)s graphed and the fits between each pump event
def pumpgrowthrates(vialOD, vialpumptimes):
    
    pumpiteration = 0
    skip = False

    if len(vialpumptimes) == 0:
        nextpumptime = float('inf')
    if len(vialpumptimes) > 0:
        nextpumptime = vialpumptimes[0]

    growth = []
    times = [0]
    growthrate = [0]
    
    nexthourtime = 2

    for x in range(len(vialOD)):
        if nextpumptime < nexthourtime+1:
            nextcalctime = nextpumptime
            
        else:
            nextcalctime = nexthourtime
            
        if np.isnan(vialOD[x][1]) == False and vialOD[x][1] > 0:
            currtime = vialOD[x][0]
            currOD = vialOD[x][1]

            if currtime < nextcalctime:
                growth.append([vialOD[x][0], vialOD[x][1]])
                
            if currtime >= nextcalctime:
                #Do fit and store data
                if len(growth) < 10:
                    skip = True
                
                if skip == True:
                    skip = False
                    
                else:
                    slope = dofit(growth)
                    times.append(currtime)
                    growthrate.append(slope)

                #Reset pumpgrowth data
                growth = []
                
                if currtime >= nextpumptime:

                    pumpiteration = pumpiteration + 1

                    if pumpiteration <= len(vialpumptimes)-1:
                        nextpumptime = vialpumptimes[pumpiteration]
                        if nextpumptime < currtime + 0.1:
                            skip = True

                    if pumpiteration > len(vialpumptimes)-1:
                        nextpumptime = float('inf')

                    
                nexthourtime = currtime + 2


            if x == len(vialOD) - 1 and len(growth) > 20:
                slope = dofit(growth)

                times.append(currtime)
                growthrate.append(slope)
    
    return (times, growthrate)

def getgenerations(times, onlypumpgrowthrate):
    generations = [0]
    for x in range(len(times)):
        if x == 0:
            continue
            
        deltat = times[x] - times[x-1]
            
        if x == 1:
            deltagen = (onlypumpgrowthrate[x+1]/ln(2)) * deltat
        else:
            deltagen = (onlypumpgrowthrate[x]/ln(2)) * deltat
            
        newgen = generations[x-1] + deltagen
        generations.append(newgen)
                     
    return(generations)
                      


#Finds the residuals from the mean of a window (default window size is 11, but can be modified by user)
#Used to find where the growth rates have stabilized (ie are not changing and thus have lower residuals from mean than if the growth rates are changing)
#Suffers when there is larger error in OD data for some reason
#Takes in the times and onlypumpgrowthrate that were made above
#Also takes in "eithersidewindow." Default is 5, calculated by taking window size, subracting 1 and then dividing by 2. Basically finding the size of the window on either side of the current value
#Returns baseindex, baseresidual, and basemean. These are the index at which the window centered around had the lowest residuals, the residual for that window, and the mean for that window.
def getwindowresiduals(times, onlypumpgrowthrate, eithersidewindow):
    
    residuals = []
    
    if len(onlypumpgrowthrate) < ((2*eithersidewindow) + 1):
        eithersidewindow = int(((len(onlypumpgrowthrate)-3) / 2))
    
    for x in range(eithersidewindow,len(onlypumpgrowthrate)-eithersidewindow):
        test = onlypumpgrowthrate[x-eithersidewindow:x+eithersidewindow+1]
        testtimes = times[x-eithersidewindow:x+eithersidewindow+1]

        mean_ = mean(test)
        deviation = 0
        for y in range(len(test)):
            rs = (test[y]-mean_)**2
            deviation = deviation + rs

        residuals.append(deviation)

    baseindex = residuals.index(min(residuals)) + eithersidewindow
    baseresidual = min(residuals) / ((eithersidewindow+1)*2)
    basemean = mean(onlypumpgrowthrate[baseindex-eithersidewindow:baseindex+eithersidewindow+1])
    
    return(baseindex, baseresidual, basemean)


#Expands the window, adding growth rate data at later time points.
#Takes in the onlypumpgrowthrate and times arrays made in pumpgrowthrates()
#Takes in baseindex, basemean, and baseresidual made in getwindowresiduals()
#Takes in residualallowed and eithersidewindow. These can be user defined, but both default to 5 on the first iteration.
#Reuturns the index relative to baseindex at which including the next growth rate would add more than "residualallowed" (5 default) times the baseresidual. Residuals are all normalized to number of data points included
#Returns index second before last relative to baseindex if it goes until the end and never exceeds residualallowed*baseresidual.
def expandright(onlypumpgrowthrate, times, baseindex, basemean, baseresidual, residualallowed, eithersidewindow):
    checkone = 0
    residuals = []
    for x in range(eithersidewindow+1,len(onlypumpgrowthrate[baseindex:])-eithersidewindow):
        test = onlypumpgrowthrate[baseindex-eithersidewindow:baseindex+x]
        testtimes = times[baseindex-eithersidewindow:baseindex+x]

        deviation = 0
        for y in range(len(test)):
            rs = (test[y]-basemean)**2
            deviation = deviation + rs

        deviation = deviation/len(test)
        residuals.append(deviation)
        if deviation > baseresidual * residualallowed:
            stop = x - 2
            checkone = 1
            break
            
    if checkone == 0:
        stop = len(onlypumpgrowthrate) - baseindex - 2
            
    return(stop)


#Expands the window, adding growth rate data at earlier time points.
#Takes in the onlypumpgrowthrate and times arrays made in pumpgrowthrates()
#Takes in baseindex, basemean, and baseresidual made in getwindowresiduals()
#Takes in residualallowed and eithersidewindow. These can be user defined, but both default to 5 on the first iteration.
#Reuturns the index relative to baseindex at which including the next growth rate would add more than "residualallowed" (5 default) times the baseresidual. Residuals are all normalized to number of data points included
#Returns index second after first relative to baseindex if it goes until the end and never exceeds residualallowed*baseresidual.
def expandleft(onlypumpgrowthrate, times, baseindex, basemean, baseresidual, residualallowed, eithersidewindow):
    checktwo = 0
    residuals = []
    for x in range(eithersidewindow+1,len(onlypumpgrowthrate[:baseindex])):
        test = onlypumpgrowthrate[baseindex-x:baseindex+eithersidewindow]
        testtimes = onlypumpgrowthrate[baseindex-x:baseindex+eithersidewindow]

        deviation = 0
        for y in range(len(test)):
            rs = (test[y]-basemean)**2
            deviation = deviation + rs

        deviation = deviation/len(test)
        residuals.append(deviation)
        if deviation > baseresidual * residualallowed:
            begin = x - 1
            checktwo = 1
            break
            
    if checktwo == 0:
        begin = baseindex - 2
    
    return(begin)


#Finds indeces corresponding with times at which the user would like the fit to occur
#Takes in the desired start and stop times (user defined)
#Takes in the times array
#Returns indeces corresponding with start and stop times in times array
def fittingbyself(start, stop, times):
    x = 0
    check = False
    while x < len(times) and check == False:
        if times[x] < start:
            startindex = copy(x) + 1
        if times[x] > stop:
            stopindex = copy(x)
            check = True
        if x == len(times) - 1:
            stopindex = copy(x)
            check = True
        x = x + 1
    
    return(startindex, stopindex)
            
        
        

###############################################################################
########################End of defining functions##############################
###############################################################################

def main():

    #Asks user to input if they are running consecutive or nonconsecutive vials
    outputfilename = input('Please enter what you would like the output file to be named (do not include .csv): ')
    os.mkdir('growth_rates')
    
    selfcheck = input("Do you want to be able to check the graphs and modify settings? (yes/no): ")
    
    vialnums, samplenames = getinfo_()

    #Initializes array that will store information about each analyzed sample
    #This will eventually be saved as a CSV file
    sampledata = []


    #Loops through each vial that was used, finding the window size, median, averages, and number of points included.
    for z in range(len(vialnums)):

        #Imports vial OD data
        vialOD = importOD(vialnums[z])

        #Imports vial pump times
        vialpumptimes = importpumptimes(vialnums[z])

        #Initializes figure on which the OD data and growth rates will be plotted on two different axes
        #ax1 contains OD data, ax2 contains growth rates
        #fig, (ax1, ax2) = plt.subplots(2, sharex = True, figsize = (10,10))
        #ax1.set_ylim([0, 0.4])
        #ax2.set_ylim([0, 1])

        #gets times of/immediately after pump events and the growth rate of the section immediately prior the pump event
        times, onlypumpgrowthrate = pumpgrowthrates(vialOD, vialpumptimes)
        
        #uses pumpgrowthrates and times to calculate generations
        generations = getgenerations(times, onlypumpgrowthrate)

        #Intializes variables, later can be user defined
        happy = "sad"
        minimumpumpsrequired = 0
        windowsize = 11
        residualallowed = 5

        #Initializes a while loop that will continue going until the user is happy with the fit
        while happy == "sad":

            #Converts the window size into how far it goes in either direction (minus the base position)
            eithersidewindow = int((windowsize - 1) / 2)

            #Gets initial window and associated values
            baseindex, baseresidual, basemean = getwindowresiduals(times, onlypumpgrowthrate, eithersidewindow)

            #Adds growth rates to window as time increases until error is too large
            stop = expandright(onlypumpgrowthrate, times, baseindex, basemean, baseresidual, residualallowed, eithersidewindow)

            #Adds growth rates to window as time decreases until error is too large
            begin = expandleft(onlypumpgrowthrate, times, baseindex, basemean, baseresidual, residualallowed, eithersidewindow)

            #Graphs the growth rates and the final window
            #ax2.scatter(times, onlypumpgrowthrate)
            #ax2.axvline(x = times[baseindex+stop])
            #ax2.axvline(x = times[baseindex-begin])

            #Stores important data in array data
            data = [samplenames[z], np.median(onlypumpgrowthrate[baseindex-begin:baseindex+stop]), np.mean(onlypumpgrowthrate[baseindex-begin:baseindex+stop]), np.std(onlypumpgrowthrate[baseindex-begin:baseindex+stop]), len(onlypumpgrowthrate[baseindex-begin:baseindex+stop]), np.median(onlypumpgrowthrate), np.mean(onlypumpgrowthrate), np.std(onlypumpgrowthrate), len(onlypumpgrowthrate), minimumpumpsrequired, windowsize, residualallowed]


            #Finishing touches on graphs and saves as a png
            #fig.suptitle(samplenames[z], fontsize = 20)
            #ax1.set(ylabel = 'OD 600')
            #ax2.set(xlabel = 'Time (h)', ylabel = 'Specific growth rate (h^-1)')
            #fig.tight_layout()
            #fig.savefig('growth_rates/vial'+str(vialnums[z])+'_'+slugify(samplenames[z])+'.png', bbox_inches = 'tight', facecolor = 'white')
                      
            if selfcheck == 'yes':
                
                from _plotly_future_ import v4_subplots
                import plotly.offline as pyo
                import plotly.graph_objs as go
                from plotly.subplots import make_subplots

                
                # Create the subplot figure
                fig_ = make_subplots(rows=3, cols=1)
                
                rawtimes = []
                rawODs = []
                for x in range(len(vialOD)):
                    rawtimes.append(vialOD[x][0])
                    rawODs.append(vialOD[x][1])

                # Add the scatter plots to the subplot figure
                fig_.add_trace(go.Scatter(x=rawtimes, y=rawODs, mode='markers'), row=1, col=1)
                fig_.add_trace(go.Scatter(x=times, y=generations, mode='markers'), row=2, col=1)
                fig_.add_trace(go.Scatter(x=times, y=onlypumpgrowthrate, mode='markers'), row=3, col=1)
                fig_.add_shape(type ="line", x0 = times[baseindex+stop], y0 = 0, x1 = times[baseindex+stop], y1 = 1, xref = 'x3', yref = 'y3', line=dict(color = "Black", width = 2, dash="dash"), row=3, col=1)
                fig_.add_shape(type ="line", x0 = times[baseindex-begin], y0 = 0, x1 = times[baseindex-begin], y1 = 1, xref = 'x3', yref = 'y3', line=dict(color = "Black", width = 2, dash="dash"), row=3, col=1)
                
                fig_.update_layout(
                    title = 'vial'+str(vialnums[z])+'_'+slugify(samplenames[z]))
                
                fig_.update_xaxes(title_text="time (h)", row = 1, col = 1)
                fig_.update_xaxes(title_text="time (h)", row = 2, col = 1)
                fig_.update_xaxes(title_text="time (h)", row = 3, col = 1)
                
                fig_.update_yaxes(title_text="OD 600", row = 1, col = 1)
                fig_.update_yaxes(title_text="Generations", row = 2, col = 1)
                fig_.update_yaxes(title_text="Growth rate (h^-1)", row = 3, col = 1)


                # Display the subplot figure
                pyo.plot(fig_, filename='growth_rates/vial'+str(vialnums[z])+'_'+slugify(samplenames[z])+'.html')

                #Checks if user is happy with the outcome, if happy will ultimately end the while loop
                happy = input('Check the figure (and leave it open or the script may fail later). Are you happy or sad with these results? (i.e., would you like to relax the analysis parameters or manually define the window) input "happy" or "sad": ')    

                #If sad, allows user to either define their own window or relax the analysis parameters
                if happy == "sad":

                    #Asks if user wants to define window themselves or relax analysis parameters
                    selffit = input('Would you like to define the window yourself, or relax the analysis parameters to find a window? (myself/computer): ')
                    if selffit == "myself":
                        while happy == "sad":
                            #Asks for user-defined window
                            start = float(input('At what time would you like to start including data in analyzing growth rate?: '))
                            stop = float(input('At what time would you like to stop including data in analyzing growth rate? If you want to include data up to the very end you can put in a time way past the last time on the graph.: '))
                            #Clears old graph and makes new graph with the user-defined window
                            #ax2.clear()

                            #ax2.scatter(times, onlypumpgrowthrate)

                            startindex, stopindex = fittingbyself(start, stop, times)
                            
                            fig_.data =[]
                            fig_ = make_subplots(rows=3, cols=1)
                            fig_.add_trace(go.Scatter(x=rawtimes, y=rawODs, mode='markers'), row=1, col=1)
                            fig_.add_trace(go.Scatter(x=times, y=generations, mode='markers'), row=2, col=1)
                            fig_.add_trace(go.Scatter(x=times, y=onlypumpgrowthrate, mode='markers'), row=3, col=1)
                            
                            fig_.add_shape(type ="line", x0 = times[stopindex], y0 = 0, x1 = times[stopindex], y1 = 1, xref = 'x3', yref = 'y3', line=dict(color = "Black", width = 2, dash="dash"), row=3, col=1)
                            fig_.add_shape(type ="line", x0 = times[startindex], y0 = 0, x1 = times[startindex], y1 = 1, xref = 'x3', yref = 'y3', line=dict(color = "Black", width = 2, dash="dash"), row=3, col=1)

                            fig_.update_layout(
                                title = 'vial'+str(vialnums[z])+'_'+slugify(samplenames[z]))

                            fig_.update_xaxes(title_text="time (h)", row = 1, col = 1)
                            fig_.update_xaxes(title_text="time (h)", row = 2, col = 1)
                            fig_.update_xaxes(title_text="time (h)", row = 3, col = 1)

                            fig_.update_yaxes(title_text="OD 600", row = 1, col = 1)
                            fig_.update_yaxes(title_text="Generations", row = 2, col = 1)
                            fig_.update_yaxes(title_text="Growth rate (h^-1)", row = 3, col = 1)


                            # Display the subplot figure
                            pyo.plot(fig_, filename='growth_rates/vial'+str(vialnums[z])+'_'+slugify(samplenames[z])+'.html')
                            

                            #ax2.axvline(x = times[startindex])
                            #ax2.axvline(x = times[stopindex])

                            data = [samplenames[z], np.median(onlypumpgrowthrate[startindex:stopindex]), np.mean(onlypumpgrowthrate[startindex:stopindex]), np.std(onlypumpgrowthrate[startindex:stopindex]), len(onlypumpgrowthrate[startindex:stopindex]), np.median(onlypumpgrowthrate), np.mean(onlypumpgrowthrate), np.std(onlypumpgrowthrate), len(onlypumpgrowthrate), "Defined by user"]

                            #fig.savefig('growth_rates/vial'+str(vialnums[z])+'_'+slugify(samplenames[z])+'.png', bbox_inches = 'tight', facecolor = 'white')

                            happy = input('Check the figure (and leave it open or the script may fail later). Are you happy or sad with these results? (i.e., would you like to relax the analysis parameters or manually define the window): input "happy" or "sad" without quotations: ') 

                    if selffit == "computer":
                        #Aks for relaxed analysis parameters.
                        minimumpumpsrequired = int(input('How many pump events do you want to be required to calculate a growth rate? (default is 10): '))
                        windowsize = int(input('What size would you like the window to be for finding the flattest region? (default is 11, number must be odd): '))
                        residualallowed = int(input('How much error would you like to allow in the residuals when the expanding the window used for calculating growth rate? (default is 5, ie 5 times the initial residuals): '))
                        fig_.data =[]
                         
            else:
                happy = "happy"

        #Adds the stored important data from data array to the sampledata array, which is eventually saved as CSV               
        sampledata.append(data)
        with open('growth_rates/'+outputfilename+'.csv', 'w', newline = '') as csvfile:
            writer = csv.writer(csvfile)
            writer.writerow(["sample name", "trimmed median", "trimmed average", "trimmed standard deviaiton", "# of cycles included", "untrimmed median", "untrimmed average", "untrimmed standard deviation", "total # of cycles in run", "Minimum pump events required", "Initial window size", "Residuals allowed"])
            for z in range(len(sampledata)):
                writer.writerow(sampledata[z])

                
if __name__ == '__main__':
    main()
